# Positional information modulates transient regeneration-activated cell states during vertebrate appendage regeneration

**DOI:** 10.1101/2024.03.28.587250

**Authors:** Augusto Ortega Granillo, Daniel Zamora, Robert R. Schnittker, Allison R. Scott, Jonathon Russell, Carolyn E. Brewster, Eric J. Ross, Daniel A. Acheampong, Kevin Ferro, Jason A. Morrison, Boris Y. Rubinstein, Anoja G. Perera, Wei Wang, Alejandro Sánchez Alvarado

## Abstract

Injury is a common occurrence in the life of organisms. Because the extent of damage cannot be predicted, injured organisms must determine how much tissue needs to be restored. It is known that amputation position determines the regeneration speed of amputated appendages in regeneration-competent animals. Yet, it is not clear how positional information is conveyed during regeneration. Here, we investigated tissue dynamics in regenerating caudal fins in the African killifish (*Nothobranchius furzeri*). We report position-specific, differential modulation of the spatial distribution, duration, and magnitude of proliferation. Regenerating fins profiled by single cell RNA sequencing identified a Transient Regeneration-Activated Cell State (TRACS) that is amplified to match a given amputation position. We located this TRACS to the basal epidermis and found them to express components and modifiers of the extracellular matrix (ECM). We propose a role for these cells in transducing positional information to the regenerating blastema by remodeling the ECM.

**Highlights:**

- Amputation position changes tissue-wide proliferation response
- Transcriptional compartmentalization is relative to injury type
- Regeneration deploys Transient Regeneration-Activated Cell States
- Prediction: positional information is transduced by ECM changes during regeneration

## Introduction

For many organisms, including humans, the preservation of anatomical form and function depends in great part on the periodic elimination and restoration of cells. This process is referred to as tissue homeostasis. In mammals, the rate of tissue homeostasis varies widely across organs; *i.e.,* self-renewal of the intestinal and lung epithelia has been estimated to take 5 days and up to 6 months, respectively^1,2^. As such, specific mechanisms exist in adult tissues responsible for sustaining specific rates of tissue homeostasis to mitigate normal, physiological wear and tear. The intricate balance of tissue homeostasis can be severely disrupted by injury. During their lifespans, all multicellular organisms are likely to experience some kind of injury. Unlike physiologically regulated tissue homeostasis, injuries and the extent of the damage incurred are unpredictable. This creates a challenge for organisms to monitor and deploy an anatomically specific regeneration response proportional to the magnitude of the injury.

During regeneration a specialized structure known as blastema forms through the rapid proliferation and subsequent differentiation of multiple cell types to restore the missing tissue^3^. For instance, it has been shown that osteoblast differentiation is accelerated in regenerating fins. Newly differentiated osteoblasts appear 4 days earlier in regenerated tissue compared to the pre-existing tissue, also referred to as the stump^4^. It is not known whether there is a regeneration-specific osteoblast differentiation program, or if differentiation trajectories used during tissue homeostasis can be accelerated during regeneration. Regeneration has also been shown to alter rates of tissue growth. In vertebrates, the speed of regeneration differs when different amounts of tissue are lost^5^. For example, fish caudal fin amputations close to the base of the appendage, a proximal amputation, display higher growth rates than amputations performed further away from the base of the fin, a distal amputation^6,7^.

To date, it is not known how injured tissues detect amputation position, and what processes may encode positional information during regeneration. Efforts to identify deposited positional information prior to injury led to the identification of transcripts and proteins that are differentially expressed along the P/D axis in the intact zebrafish fin^8^. However, these differences are lost during regeneration leading to the hypothesis that positional information must be redefined during regeneration^8^. Alternatively, it has been proposed that migratory progenitor cells retain positional identity from their locations prior to injury. This opens up the possibility that progenitors communicate positional information to the new tissue to determine regeneration growth rate^9^. But the mechanism of positional information retention and potential relay to other cells has not yet been identified.

Furthermore, it has been suggested that positional information is encoded in the tissue within the thickness of the bone at the plane of amputation^10,11^, and that mechanical distension of the epidermis during wound closure constitutes a direct measurement of the amputation position by the wounding epidermis. A wave of mechanical distension in the basal epidermis was shown to propagate to different lengths according to the amputation position^12^. To add to the complexity of this process, it was shown that there is a 2-day time window at the beginning of regeneration when positional information is possibly reestablished *de novo*. If blastemas are impaired in their proliferative ability during this time window, the wrong positional information is encoded in the regenerated appendage, so multiple rounds of regeneration consistently grow abnormal tissue sizes in the absence of any further impairment of proliferation^13^. It is conceivable that independent mechanisms of positional information coordinating the regenerative response exist. There may be redundant and complementary means to adequately relay and deposit positional information into the new tissue. This would ensure that form and function are restored following diverse and unpredictable injury.

Here we deploy spatial and temporal analysis of proliferation, and single cell transcriptomic profiling to measure molecular and cellular changes along the caudal fin P/D axis. We chose the African killifish *Nothobranchius furzeri* for our studies because of the reduced complexity of differential gene expression and gene regulation compared to the more broadly utilized zebrafish^14^. We show that amputation position influences the length of time it takes for tissues to progress through regeneration. We report on the discovery of a basal epidermal subpopulation that shows a Transient Regeneration-Activated Cell State (TRACS) likely to participate in mediating positional information in the regenerating blastema. Altogether, our study demonstrates that amputations along the P/D axis result in defined spatial and temporal rates of proliferation. We propose that such dynamics likely initiate the proportional changes in tissue architecture that may ultimately define the scale and rate of regeneration of amputated tissues.

## Results

### Amputation position influences regeneration growthrate in killifish caudal fins

We investigated whether the killifish *N. furzeri* displays regeneration growth rate differences at different amputation positions in the caudal fin. We restricted our analyses to male individuals of the same size because the caudal fin has a clear pigmentation pattern that allows for consistent amputation at the same position across multiple individuals (Figure 1A, Supplementary Figure S1A, S1C, S1D). Whether there are sex-determined differences that influence regeneration growth rate in killifish remains to be investigated. We first established the effect of amputation position by carrying out regeneration time courses at two different amputation positions along the P/D axis (Figure 1B, Supplementary Figure S1E). We quantified growth rate (mm/day) by measuring the regeneration length and dividing by the time between adjacent time points (Figure 1D). Consistent with other fishes, killifish showed regeneration growth rate differences when comparing distinct amputation positions, with proximal injuries displaying a higher growth rate compared to distal injuries (Figure 1C, Supplementary Figure S1B). We observed growth rate differences between proximal and distal injuries within 2- and 7- days post amputation (dpa), with the peak growth rate difference occurring at 3.5 dpa, and both injuries reaching and maintaining the same growth rate from 7 dpa onward (Figure 1E-F). We also observed significant differences in growth acceleration and deceleration when comparing both proximal and distal injuries (Figure 1G).

**Figure 1.**
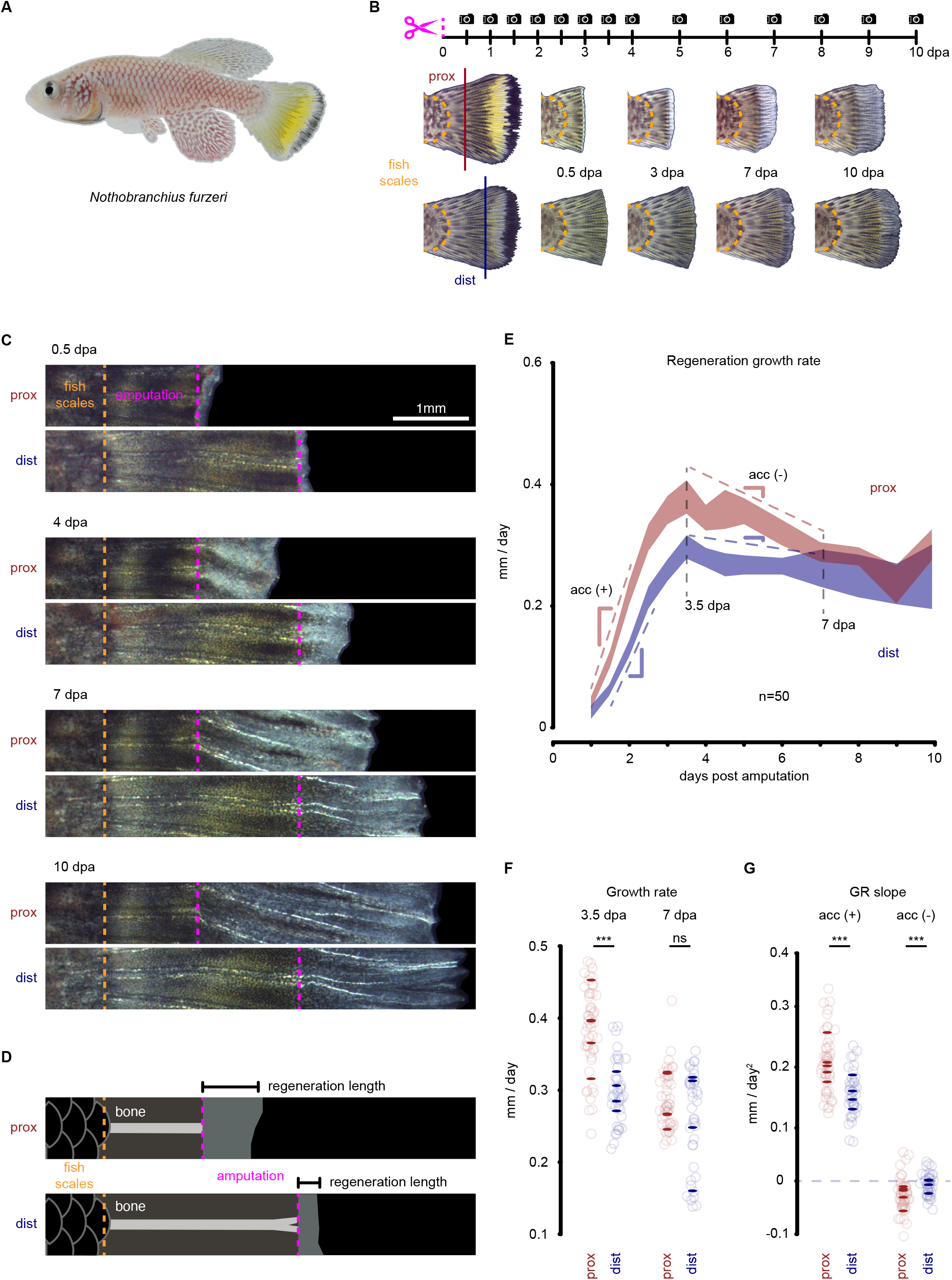
Amputation position triggers differential regenerative outgrowth in killifish. (A) Sideview of sexually mature adult male killifish. (B) Experimental design for longitudinal analysis of regenerative outgrowth. (C) Representative images of regeneration time course of proximal and distal injuries aligned at the boundary between the fish scales and the caudal fin (orange dotted line), amputation line is displayed in magenta, scale bar 1 mm. (D) Definition of regeneration length in the context of the new tissue, pre-existing tissue, bone, amputation, and fish scales. The difference in regeneration length divided by the time between time points corresponds to the regeneration growth rate expressed in mm/day. (E) Regeneration growth rate in proximal and distal injuries. Grey dashed vertical lines correspond to the most different growth rate or the earliest time point when growth rate is the same between proximal and distal injuries respectively. Colored dashed lines with indicated slopes correspond to growth acceleration and slowing down (n=50). (F) Statistical analysis of growth rate quantified at 3.5 and 7 dpa. Open circles reflect individual bones, filled ovals correspond to individual fish. (G) Statistical analysis of growth acceleration and slowing down. Open circles and filled ovals represent the same as in (E). ∗∗∗ *FDR<0.001*, Wilcoxon rank sum test. See also Supplementary Figure S1.

We conclude from these experiments that regenerative outgrowths have at least two components that are influenced by amputation position: 1) a magnitude component, as indicated by the higher growth rates of proximal injuries when compared to distal injuries (Figure 1E, 1F); and 2) a time component in which proximal injuries progress to higher growth rates at a sooner time point compared to distal injuries (Figure 1E, 1G). This later observation may not have been detected in prior studies due to the lower temporal resolution reported^7,10^ (Supplementary Figure S1B). We reasoned that the first component could be explained by expanding the progenitor pool to match the amputation position as previously observed^7,11^. However, the second component suggested an additional layer of regulation that influences the pace at which progenitors become activated to mount a corresponding regenerative response to the amputation position. We hypothesized that amputation position changes both the number of progenitor cells recruited to the regeneration event, and the amount of time pro- proliferative signals persist in the tissue. We tested this idea by quantifying the cellular proliferation dynamics in proximal and distal injuries during the formation of the initial stages of blastema formation and before the tissue undergoes regenerative outgrowth: 12 to 48 hours post amputation (hpa).

### Amputation position determines the duration of tissue-wide proliferation

We previously reported that between 14 and 24 hpa fins display high levels of proliferation in the pre-existing tissue (tissue-wide proliferation), while earlier time points have proliferation levels similar to unamputated fins, and later time points display high levels of proliferation localized to the plane of amputation (blastema proliferation)^14^. We investigated the effect of amputation position on tissue-wide proliferation over time by whole-mount immunofluorescence against the mitosis marker phospho-Histone H3 (H3P), automatic nuclei segmentation, and cytometry analyses (Figure 2A-C, Supplementary Figure S2A). We observed that both proximal and distal injuries displayed tissue-wide proliferation at 18 hpa (Figure 2C, 18 hpa prox dist). However, the distribution of proliferative events diverged after 18 hpa, with distally amputated fins reducing cell division in fin tissue located away from the plane of amputation (-1.5 mm), and proximally amputated fins sustaining such proliferation for an additional 18 hours (Figure 2C’, asterisks). In contrast, the tissue adjacent to the plane of amputation (-0.075 mm) in both distally and proximally amputated fins displayed a sustained increase in proliferation (Figure 2C”). We also observed a positive correlation between increased proliferation events in thicker (proximal) than in thinner fin tissues (distal) as previously described^7^ (Supplementary Figure S2B-D). Thus, amputation position appears to influence the length of time the regenerating tissue undergoes tissue-wide proliferation (Figure 2D). We conclude from these experiments that the number of cells recruited for cell division as well as the duration of tissue-wide proliferation depend on the location of the amputation plane and that both events likely determine blastema size and regeneration growth rate.

**Figure 2.**
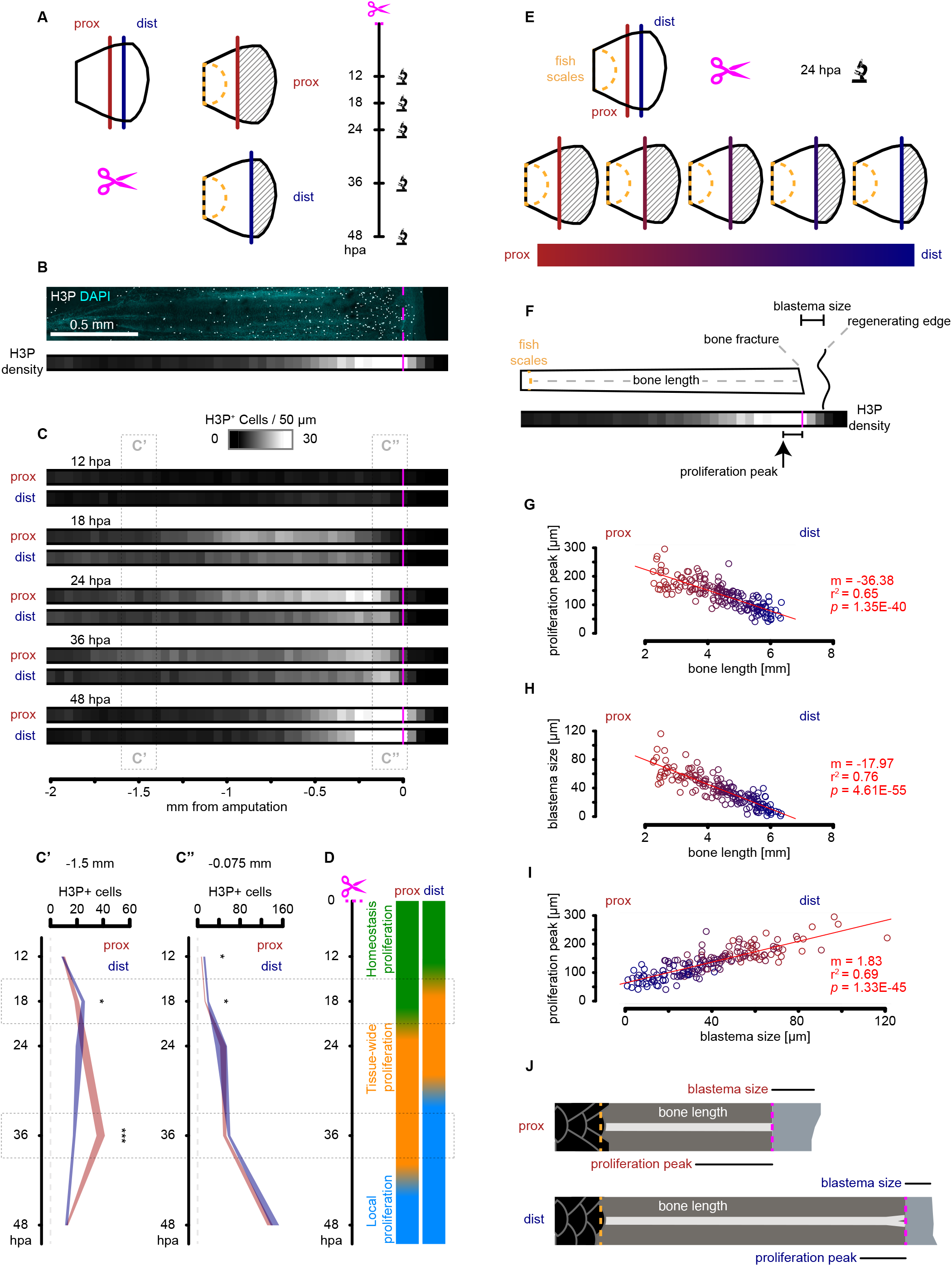
Amputation position influences temporal and spatial distribution of proliferating cells. (A) Experimental design of regeneration time course at two different amputation positions. (B) Top: Single bone representative image of masked H3P in greyscale with cyan DAPI background, scale bar 0.5 mm. Bottom: heatmap representation of H3P density along the P-D axis relative to the amputation plane (C) Heatmap representation of averaged H3P density in regenerating fins at different amputation positions. Time course is displayed in vertical orientation, n=12. (C’) Statistical analysis per time point corresponding to 200 μm segment 1.5 mm away from the amputation plane, polygon area represents mean ± sem, n=12. (C’’) Statistical analysis per time point corresponding to 200 μm segment adjacent to the amputation plane, polygon area represents mean ± sem, n=12. (D) Regeneration progression model shows proliferation time shift between proximal and distal injuries where tissue-wide proliferation is launched sooner in distal injuries, and it is maintained for longer in proximal injuries. (E) Experimental design of gradient-cut regeneration dataset where images were collected at 24 hpa. (F) Blastema size and proliferation peak definition relative to bone length, bone fracture, and regenerating edge. (G) Linear correlation between proliferation peak distance to the amputation and bone length. Larger bones correspond to distal amputations. (H) Linear correlation between blastema size and bone length. (I) Linear scaling of 1.83 μm proliferation peak distance to the amputation per 1 μm of blastema size. (J) Schematic of length relationship between blastema size and proliferation peak in proximal and distal injuries.

### Amputation position predicts blastema size and spatial distribution of proliferation

Interestingly, most dividing cells during tissue-wide proliferation (24 hpa) are located within the epidermis (Figure S2A). We reasoned that amputation position may shape the spatial distribution of tissue-wide proliferation. To test this idea, we generated spatial proliferation profiles in a proximal-distal gradient of amputations. We performed immunostaining against H3P in a cohort of fish amputated at different positions along the P/D axis and quantified the resulting proliferation profiles at 24 hpa (Figure 2E). Previous work has shown that proximal injuries have greater numbers of dividing progenitor cells than distal injuries^7^. We reasoned that the spatial distribution of dividing cells may reflect an interplay between how many cells are dividing and for how long they are being deployed tissue-wide. We investigated the possibility of a spatial control mechanism by calculating the distance between the proliferation peak, corresponding to the highest density of proliferating cells, with the plane of amputation (Figure 2F). We observed that bone length, which is a proxy for amputation position, significantly correlates with both the proliferation peak distance to the plane of amputation (*p<0.001*, Figure 2G) and the size of the blastema (*p<0.001*, Figure 2H). At 24 hpa, blastemas are mainly composed of wounding epidermis, which migrates to cover the exposed tissue after amputation^15,16^. We observed that proximal injuries have larger blastemas than distal injuries, which is consistent with the displacement of more epidermis to close a larger wound. The scale of epidermis displacement may propagate information further away from the injury into the pre-existing tissue which is what we observed when correlating blastema size with proliferation peak (Figure 2I). This positive correlation (by a factor of 1.83) suggests a proportional spatial remodeling of the pre-existing tissue that matches the amputation position (Figure 2J). We hypothesize from these findings that a cellular entity may exist that relays such positional information and sets the regeneration event to recruit migrating mesenchyme for a correct blastema size and growth rate. To test this hypothesis, we profiled the pre-existing tissue by single cell RNA sequencing (scRNAseq) at 12 hpa, right before the tissue-wide proliferation response is observed.

### Single cell expression analyses reveal transcriptional diversity in intact and regenerating fins

We sought to determine whether cellular and gene expression differences could be identified along the P/D axis of the regenerating caudal fin. We chose to use single cell transcriptomic profiling because of the inconclusive differences between proximal and distal injuries observed in published bulk measurements^8^. We generated three scRNAseq regeneration datasets using the 10x Genomics CellPlex multiplexed technology that allowed us to both increase the number of cells analyzed and to generate biological replicates per sample (Supplementary Figure S3A, see methods). First, we sampled regenerating and control fins sharing equivalent anatomical compositions (caudal dataset, Figure 3A). Second, we collected intact dorsal fins from unamputated control fish (dorsal homeostasis, Figure 3E) and intact dorsal fins from fish regenerating their caudal fin for 12 hours (dorsal 12 hpa, Figure 3E). And third, we profiled 12 hpa stumps from fins with proximal or distal injuries (Figure 3I). This strategy allowed us to distinguish the transcriptional responses resulting from amputation position, tissue anatomy, and regeneration.

**Figure 3.**
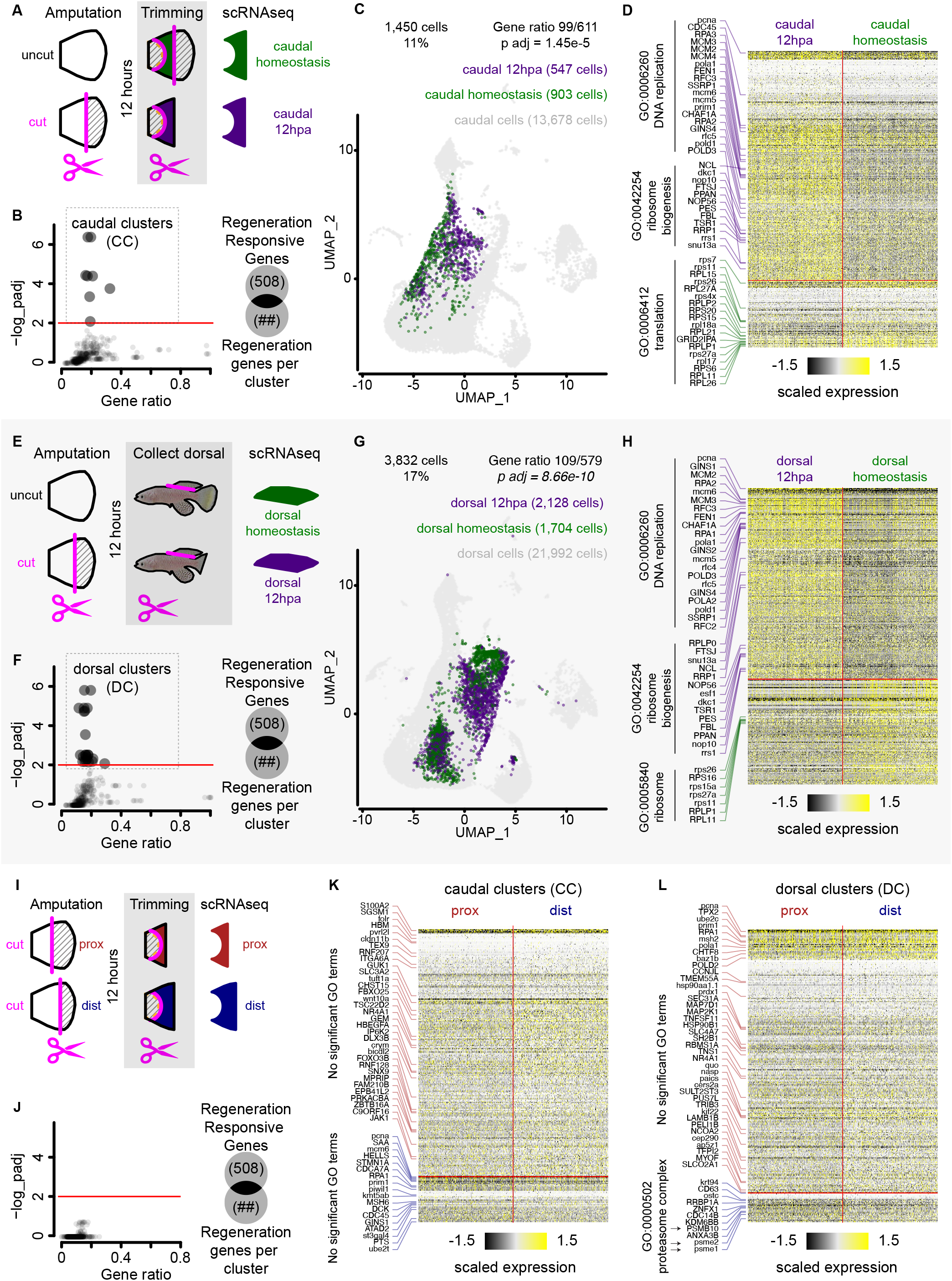
Amputation induces regeneration tran scriptional response within epidermis subpopulation at 12 hpa. (A) Experimental design of 12 hpa and homeostasis tissue collection for scRNAseq on the caudal fin. Fins were uncut or 50% cut, 12 hours later the stump was trimmed to match the corresponding anatomical region between the two samples, then tissues were dissociated and subjected to scRNAseq. (B) RRG enrichment analysis on DE genes between 12 hpa and homeostasis caudal cells split by clusters defined in the integrated dataset (656 clusters), Fisher’s exact test, *FDR < 0.01*. (C) UMAP distribution of caudal clusters (CC) within the caudal dataset, cell numbers per sample, and RRG enrichment analysis, Fisher’s exact test, *p < 0.001*. (D) Heatmap of DE genes between caudal 12 hpa and caudal homeostasis, and top GO terms associated with the gene list (*FDR < 0.05*). Cells were randomly sampled to balance both 12 hpa and homeostasis groups. (E) Experimental design of 12 hpa and homeostasis intact dorsal fin collection for scRNAseq. Caudal fins were uncut or 50% cut, 12 hours later the intact dorsal fin was collected, dissociated, and subjected to scRNAseq. (F) RRG enrichment analysis on DE genes between 12 hpa and homeostasis dorsal cells split by clusters defined in the integrated dataset (656 clusters), Fisher’s exact test, *FDR < 0.01*. (G) UMAP distribution of dorsal clusters (DC) within the dorsal dataset, cell numbers per sample, and RRG enrichment analysis, Fisher’s exact test, *p < 0.001*. (H) Heatmap of DE genes between dorsal 12 hpa and dorsal homeostasis, and top GO terms associated with the gene list (*FDR < 0.05*). Cells were randomly sampled to balance both 12 hpa and homeostasis groups. (I) Experimental design of proximal and distal regenerating caudal fins collected for scRNAseq. Fins were amputated either proximal or distal, 12 hours later the stump was trimmed to remove the scales and muscle, then tissues were dissociated and subjected to scRNAseq. (J) RRG enrichment analysis on DE genes between proximal and distal cells split by clusters defined in the integrated dataset (656 clusters), Fisher’s exact test, *FDR < 0.01*. (K) Heatmap of DE expressed genes between proximal and distal cells within the caudal clusters (CC). No significant GO terms were found (*FDR < 0.05*). Cells were randomly sampled to balance both proximal and distal groups. (L) Heatmap of DE expressed genes between proximal and distal cells within the dorsal clusters (DC), and single GO terms associated with the gene list (*FDR < 0.05*). Cells were randomly sampled to balance both proximal and distal groups.

We then integrated the obtained multiplexed scRNAseq datasets above alongside a previously published blastema dataset^14^ to generate gene expression clusters and a cell identity reference atlas (see methods for integration and quality control). The resulting integrated dataset contained 84,616 cells distributed into 32 cell clusters (cluster resolution: 1), belonging to 11 cell types expected to reside in fin tissues (Supplementary Figure S3B). We assigned cell clusters to cell types based on both known molecular markers and gene ontology enrichment analysis (Supplementary Figure S3C). To identify any cell states that may be regeneration-activated in a larger array of potential cell states, we generated a collection of 656 clusters by iteratively changing the cluster resolution in the integrated dataset (cluster resolution: 0.2-2). By doing this, we generated clusters that partially overlapped with each other and represented different degrees of transcriptional variance. This approach allowed us to leverage the transcriptional diversity captured across multiple experiments with cells obtained specifically from cut (regeneration) and uncut (homeostasis) caudal and dorsal fins.

### Epidermis subpopulation expresses early regeneration-responsive transcriptional signature

To generate differentially expressed (DE) gene lists, we split the integrated dataset into caudal and dorsal datasets (13,678 and 21,992 cells respectively) and compared cells from homeostasis and regeneration samples to each other. In the caudal fin, out of 656 clusters, 337 clusters (51%) showed more than 10 DE genes, and 219 clusters (33%) showed more than 50 DE genes. Similarly, in the dorsal fin 353 clusters (54%) showed more than 10 DE genes, and 279 clusters (42%) showed more than 50 DE genes. We asked whether the DE gene lists obtained for each cluster were significantly enriched for previously reported Regeneration Responsive Genes (RRG)^14^. We found 8 caudal clusters (CC, Figure 3B), and 23 dorsal clusters (DC, Figure 3F) with statistically significant enrichment for RRG (*padj<0.001*). Interestingly, all RRG enriched clusters belonged to the epidermis. We investigated the degree of overlap between the CC and DC by going back to the integrated dataset and grouping together all cells within each definition. We observed that 86% of cells within the CC and 88% of cells within the DC are not present within the reciprocal definition and concluded that both CC and DC represent different epidermal cell states (Figure 3C, 3G). We considered the possibility that data integration failed and that the high degree of non-overlap between CC and DC were batch effect artifacts. However, we confirmed the presence of dorsal cells within the CC and caudal cells within the DC (Supplementary Figure S3D). We interpret these data to mean that the identified CC and DC represent epidermal cell states that respond to regeneration in a context-dependent manner.

We next identified DE genes between regeneration and homeostasis in both caudal and dorsal cells (1,450 and 5,259 cells, respectively) within the CC, and caudal and dorsal cells (2,558 and 3,832 cells, respectively) within the DC (Supplementary Figure S3D). Interestingly, half the cells within the CC belong to the proliferating epidermis as indicated by elevated expression levels of proliferation markers (*e.g.*, *ccna*, *mki67* and *pcna*; Supplementary Figure S3E). We also confirmed a highly significant overrepresentation of RRG in all four cell groups albeit exclusively within the upregulated component (*padj<0.001*, Supplementary Figure S3F). Additionally, we observed a high degree of overlap (∼40%) of DE genes within all four groups of cells suggesting that the two epidermal subpopulations appear to be mounting a similar transcriptional response to regeneration.

To further investigate the transcriptional signature observed in CC and DC, we performed gene ontology (GO) term analysis on the differentially expressed gene list (Figure 3D, 3H; Supplementary Figure 3G and Supplementary Table). We found 110 and 115 GO terms in CC and DC, respectively. Within the upregulated genes, top terms (*padj<0.05*) were represented by DNA replication (GO:0006260) and ribosome biogenesis (GO:0042254). Conversely, within downregulated genes, the most significant terms were associated with translation (GO:0006412) and ribosome (GO:0005840). Interestingly, we found 144 regeneration responsive genes with a CC bias enriched for unfolded protein binding (GO:0051082), 180 genes with a DC bias enriched for genes involved in DNA replication (GO:0006260) and DNA repair (GO:0006281), and 250 genes with no cluster bias with genes enriched for the proteasome complex (GO:0000502) and protein catabolic process (GO:0030163, Supplementary Figure S3G). We conclude from our analyses that injury triggers a local and remote RRG transcriptional response in epithelial cells of both cut (caudal) and uncut (dorsal) fins. In uncut dorsal fins the response is detected in cells already committed to proliferation (dorsal clusters, Figure 3F), while in cut fins, the response is found in a subtype of non-proliferative cells (caudal clusters, Figure 3C).

### Amputation position biases differential cell state abundance rather than differential gene expression

In the search for the effects of amputation position on the deployment of regeneration, we profiled 12 hpa stumps from fins with proximal or distal injuries with scRNAseq (Figure 3I). Because proximal and distal injuries generate stumps with slightly different anatomies, we needed first to understand the transcriptional response to regeneration (see previous section). Similar to what we described above, we looked for DE genes between proximal and distal cells in all 656 clusters defined in the integrated dataset. Within these, we found 577 clusters (88%) with more than 10 DE genes and 552 clusters (84%) with more than 50 DE genes. However, we found zero cluster identities with an enriched regeneration-responsive transcriptional signature when we overlap the DE gene lists with the RRG (Figure 3J). This is not surprising since both samples correspond to the same regeneration time point that is mounting a regenerative response. This is more apparent when we look for the DE genes within the epithelial subpopulations identified above. In the CC, we observed 220 genes upregulated in proximal cells and 41 genes upregulated in distal cells (Figure 3K; Supplementary Table). When we look at the DE genes within the DC, we observe 248 genes upregulated in proximal cells and 28 genes upregulated in distal cells (Figure 3L; Supplementary Table). Surprisingly, we could not find any significant enrichment for GO terms among the DE expressed genes in both cluster groups except for the smallest gene list corresponding to 28 upregulated genes in distal cells of the DC (*padj<0.05*). The GO terms correspond to proteosome complex, peptidase complex, endopeptidase complex, catalytic complex (GO:0000502, GO:1905368, GO:1905369 and GO:1902494 respectively) and they were all part of the regeneration transcriptional signature present in both CC (Figure 3B) and DC (Figure 3F, Supplementary Figure S3G). Furthermore, we investigated whether CC or DC show an enrichment for RRG and found that genes upregulated in distal cells within CC have a statistical enrichment for RRG (*padj<0.001*, Supplementary Figure S3F). The distal bias in regenerative gene expression may be interpreted as a temporal bias meaning that regeneration is deployed sooner in distal injuries thus shortening the time window for blastema formation. In accordance with previous observations^8^, our data confirms that both proximal and distal injuries launch a transcriptionally equivalent response during regeneration. Hence, we next investigated whether amputation position influenced the relative abundance of cell states to look for a differentiation bias between proximal and distal samples.

An alternative to changes in gene expression is that amputation position may favor a particular cell cluster to numerically expand within the tissue during regeneration, thus resulting in differential abundance of a given cell cluster. This kind of transcriptional output would be the result of amputation position favoring differentiation into specific cellular states. To investigate this possibility, we performed differential abundance analysis using the bioinformatic tool Milo^17^ that computes differential abundance between two samples using *k*-nearest neighbor graphs and tests the statistical enrichment of cells for a given sample to partially overlapping neighborhoods (Figure 4A). We observed that some neighborhoods in all major cell types in the fin show statistically significant differential abundance between proximal and distal samples (*FDR<0.05*). However, two cellular identities showed a higher proportion of differentially abundant neighborhoods: basal epidermis 2 (bsEp2) and mesenchymal 4 (mes4; Figure 4B). We further investigated the differential abundance bias in our caudal dataset where both homeostasis and regeneration samples correspond to equivalent anatomical regions of the tissue (Figure 3A). We found that bsEp2 is significantly more abundant during regeneration compared to homeostasis with all neighborhoods associated to this cluster identity showing significant abundance enrichment during regeneration (*FDR<0.05;* Figure 4C). We reasoned that the overlap between the two differential abundance analyses reflect a cell differentiation bias influenced by both regeneration and amputation position. Moreover, we were able to filter out expected abundance biases such as the one observed for mes4 because there is no neighborhood within this cluster that displays abundance enrichment between regeneration and homeostasis (*FDR<0.05;* Figure 4C). We believe the abundance differences captured by the proximal and distal comparison in the mes4 cluster are likely due to intrinsic morphological differences between both injuries that are not present in the regeneration and homeostasis comparison.

**Figure 4.**
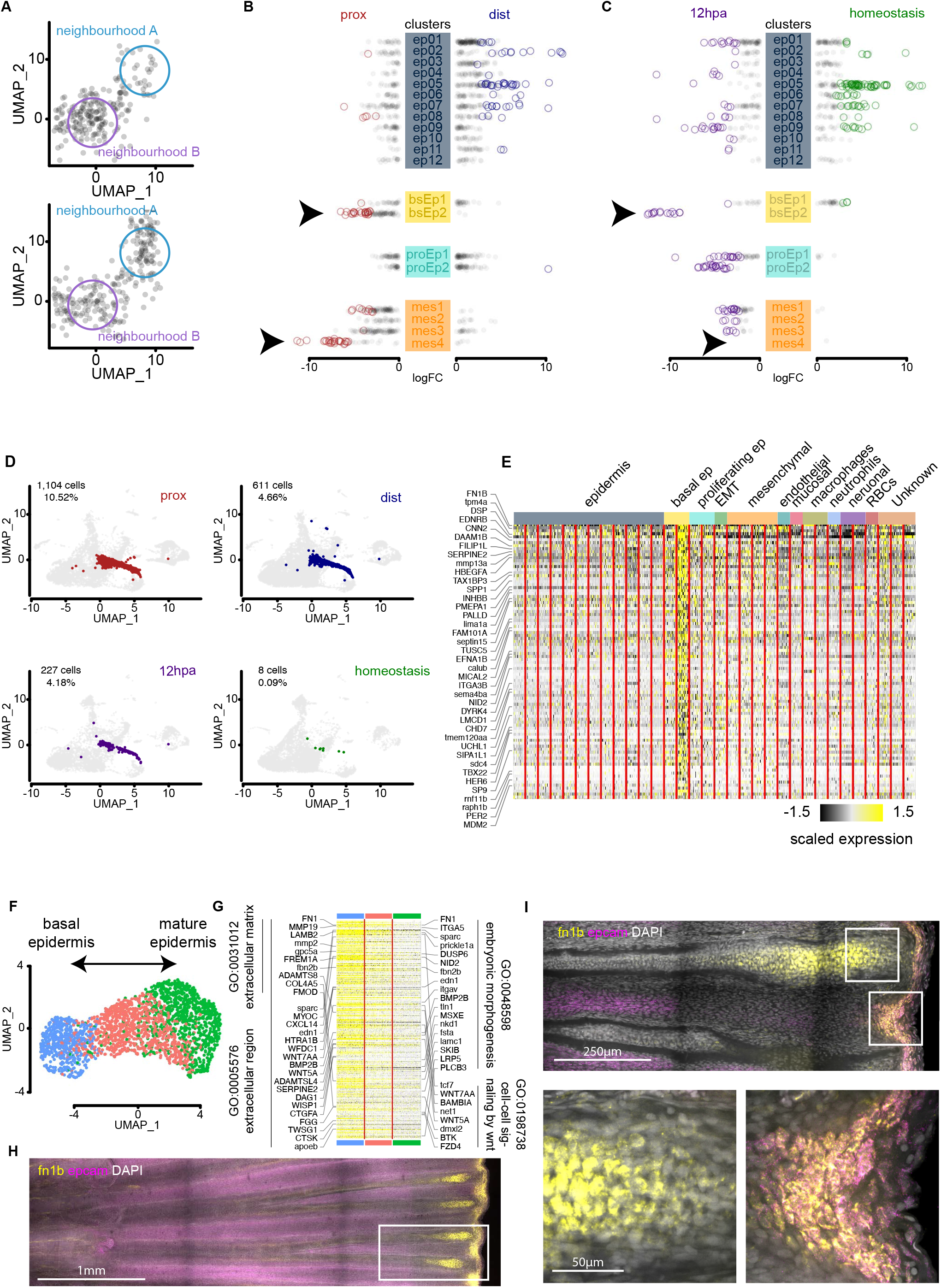
Differential abundance analysis identifies *fn1b^+^ epcam^+^* regeneration-activated basal epidermis subpopulation. (A) Two-dimensional representation of differential abundance (DA) example analysis where two neighborhoods (colored circles) surround cells are depicted with black. Note that the algorithm performs the neighborhood analysis in a K-nearest neighborhood (KNN) graph. (B) DA analysis of proximal and distal samples with statistically significant neighborhoods represented in color, the four most abundant cell types are displayed with the cluster definition from Supplementary Figure S3. (*FDR < 0.05*), arrowheads correspond to basal epidermis 2 (bsEp2) and mesenchymal 4 (mes4) clusters. (C) DA analysis of 12 hpa and homeostasis samples with statistically significant neighborhoods represented in color, displayed cell types, statistical significance and arrowheads is the same as in (B). (D) Dimensional reduction UMAP plots of regeneration-activated basal epidermis cell state across proximal, distal, 12 hpa and homeostasis samples. In color is depicted the bsEp2 cluster on top of a background with the remaining cells of the sample. (E) Heatmap of marker genes unique to the bsEp2 against all other clusters in the integrated dataset. No GO terms were found to be statistically enriched in this gene list (*FDR<0.05*). (F) Dimensional reduction UMAP distribution of subcluster analysis of bsEp2. (G) Heatmap of downregulated genes that are common in the potential differentiation trajectory from basal epidermis to epidermis, alongside significant GO terms (*FDR < 0.01)*. (H) MAX projection of whole-mount HCR *in situ* hybridization of 1 dpa regenerating tail fin. *fn1b* is displayed in yellow, *epcam* in magenta, and DAPI in greyscale. White box corresponds to top image in panel (I), scale bar 1 mm. (I) 10 μm MAX projection of the same sample depicted in panel (H) imaged at higher resolution. White squares correspond to *fn1b*+ expression domains within the tissue. Bottom left corresponds to mesenchymal cells housed within the bone plates of the fin. Bottom right is regeneration-activated basal epidermis (bsEp2) co-expressing the markers *fn1b* and *epcam*.

We further investigated the role of bsEp2 (Figure 4B, arrow) in transducing information of amputation position into the regenerating tissue. We observed that in the integrated UMAP space (Supplemental Figure 3A), different to most other cell populations, bsEp2 displays a linear trajectory along the UMAP space in between basal epidermis 1 (bsEp1) on one end, and proliferating epidermis 1 (proEp1) and epidermis 5 (ep05) populations on the other end (Supplemental Figure S3B). We looked at the possibility that the differential abundance present between proximal and distal injuries comes from a delay in differentiation displayed as a bias towards one end of the differentiation trajectory. However, in both injuries the bsEp2 cells are evenly distributed along the UMAP trajectory and it is only their numbers that differ between proximal and distal samples (Figure 4D). We looked for marker genes that distinguish bsEp2 from the rest of the cell clusters and found secreted molecules and modifiers of the extracellular space: *fn1b*, *mmp13a*, *serpine2* and *hbegfa*. We also found genes responsible for cytoskeleton structure or remodeling: *tpm4a*, *dapk3*, *plek2*, *lima1a* and *myo9b* (Figure 4E; Supplementary Table). Altogether, our data indicate that differential abundance of cell states, rather than differential gene expression, are the most detectable changes during regeneration at different amputation positions along the P/D axis of the caudal fin.

### Regeneration-activated basal epidermis remodels the ECM at the injury site

To better understand the transcriptional output of the regeneration-activated bsEp2 subpopulation, we isolated these cells and redefined the PCA and UMAP embedding configurations. We observed three continuous subtypes of regeneration-activated basal epidermis that we interpret to be a differentiation trajectory between basal epidermis and mature epidermis (Figure 4F). We performed differential expression analysis alongside the differentiation trajectory presuming a fate transition from basal epidermis into mature epidermis. We found that 13% and 24% of DE genes were upregulated, while the corresponding 87% and 76% of DE genes were downregulated alongside the differentiation trajectory. The data are consistent with previous observations that regeneration promptly shuts down injury activated transcriptional programs to shift the microenvironment from a scarring-inducing injury to a regeneration competent event^14,18,19^. We further investigated the genes that are being downregulated across the regeneration-activated basal epidermis differentiation trajectory. We observed a significant enrichment for genes that are part of the extracellular matrix (GO:0031012), localize to the extracellular region (GO:0005576), participate during embryonic morphogenesis (GO:0048598), and mediate cell-cell signaling by Wnt (GO:0198738, Figure 4G; Supplementary Table). Animals deficient in interleukin-11 signaling display impaired regeneration because of aberrant deposition of fibronectin in the regenerating hearts and poor mesenchymal migration during fin regeneration^20^. Interestingly, we found a significant enrichment of the genes downregulated along the bsEp2 differentiation trajectory within the downregulated orthologues in the *il11ra* knock-out mutant (*padj<0.001*). This data predicts a deficiency in regeneration-activated basal epidermis differentiation that leads to what the authors defined as mammalian-like fibrosis and failure to regenerate^20^. We propose that the regeneration-activated basal epidermis at the plane of amputation undergoes mechanical distension during wound healing, and upregulates genes know to remodel the cytoskeleton and ECM. Similar ECM remodeling programs during regeneration have been observed in mammalian fibroblast during skin ^21^⍰.

We used confocal microscopy to visualize the regeneration-activated basal epidermis at 1 dpa using *in situ* hybridization chain reaction (HCR^22^) probes targeting Epithelial Cell Adhesion Molecule (*epcam*, ENSNFUG00015018342) and Fibronectin 1b (*fn1b*, ENSNFUG00015003666). We observed a robust upregulation of *fn1b* adjacent to the plane of amputation (Figure 4H). Interestingly, we observed two domains of *fn1b* expression that correspond to different cell populations 1) *epcam*^+^ *fn1b* ^+^ cells localized at the interface between the wounding epidermis and the exposed mesenchyme, and 2) *epcam*^-^ *fn1b* ^+^ cells localized inside of the bone one segment away from the plane of amputation (Figure 4I). This result made us return to look for *fn1b*^+^ mesenchymal cells in our integrated dataset, we found 617 cells scattered across all four mesenchymal clusters that accounts for 7% of the mesenchymal compartment. Because they do not separate from the main mesenchymal clusters, we conclude that there are not enough *fn1b*^+^ mesenchymal cells captured in our experiments to further investigate a regeneration-activated mesenchymal cell state.

### The regeneration-activated *fn1b*^+^ basal epidermis is a transient cell state

We investigated the spatial and temporal distribution of *fn1b*^+^ basal epithelial cells in proximal and distal amputations by HCR *in situ* hybridization and antibody stains against E-cadherin (Ecad, a.k.a. Cdh1) and H3P. We observed *fn1b* expression in epithelial cells as soon as 3 hpa, localized to the amputation plane in cells that accumulate at the wound epidermis. At 3 hpa both proximal and distal injuries show *fn1b* expression. Proximal injuries show *fn1b*^-^ cells intermingling with *fn1b*^+^ epithelial cells. In contrast, distal injuries show a homogeneous population of epithelial cells expressing *fn1b* at the amputation plane (Figure 5A). At 6 hpa there is an increase in *fn1b*^+^ expressing cells, and transcript localization shifted from nuclear to cytoplasmic. Both proximal and distal injuries display strong *fn1b* expression in multiple cell layers at the wounding epidermis (Figure 5B). At 1 dpa, *fn1b* expression continues to localize to multiple cell layers but distal injuries display a reduced expression domain with sharper boundaries between the underlying mesenchyme and the most superficial *fn1b*^-^ epidermal cell layers (Figure 5C). At 2 dpa, both proximal and distal injuries show narrow *fn1b* expression domains unambiguously labeling the boundary between the migrating mesenchymal blastema and the wounding epidermis (Figure 5D). Moreover at 3 and 5 dpa, *fn1b* expression is shut down (Supplementary Figure S4A-B), indicating that the basal epidermal population that upregulates *fn1b* upon injury may be a transient regeneration-activated cell state (TRACS).

**Figure 5.**
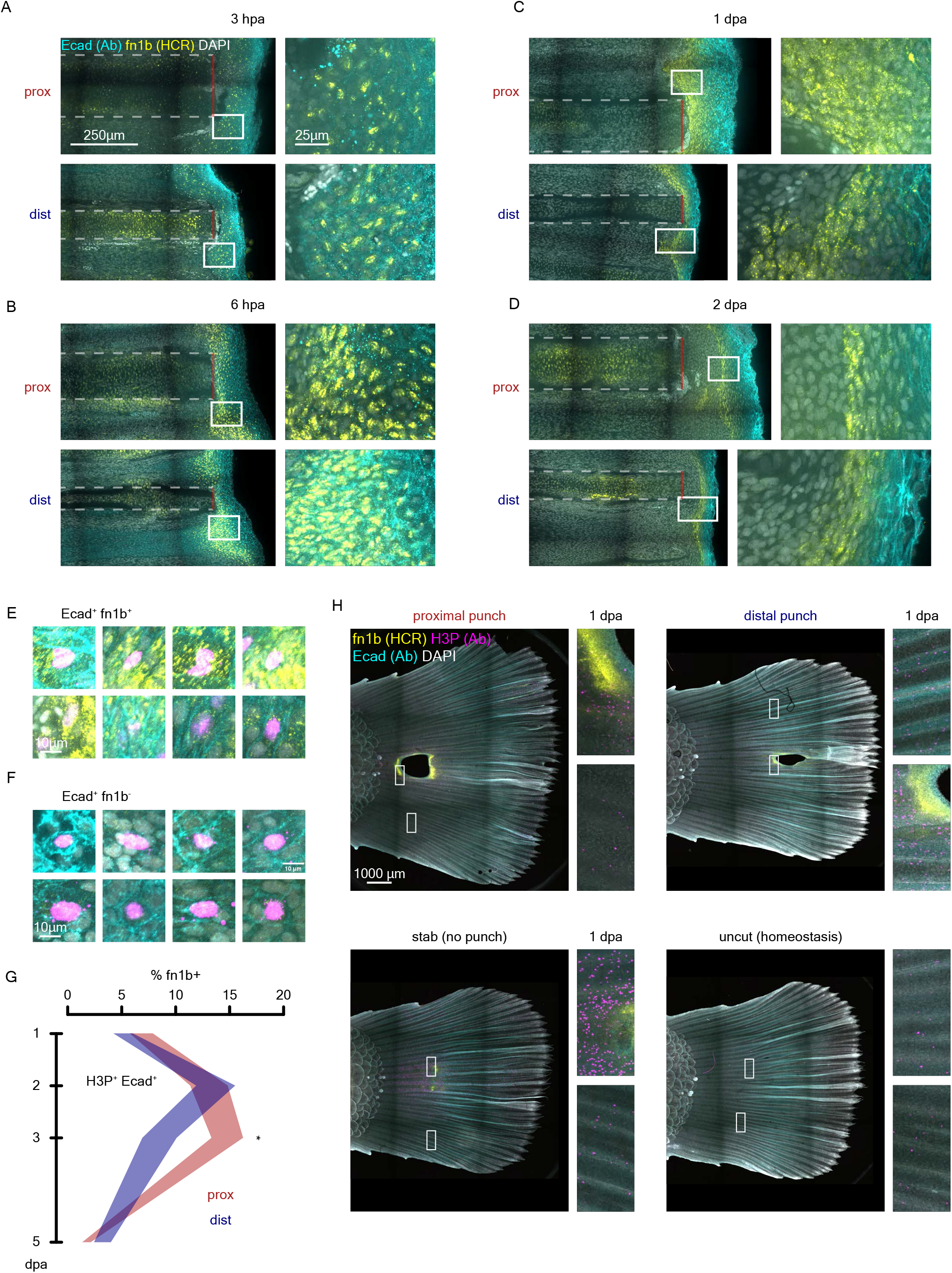
Regeneration induced *fn1b*^+^ Ecad^+^ basal epithelial subpopulation is transient and non-proliferative. (A) Early nuclear *fn1b* expression detected by HCR *in situ* hybridization in wounding epidermis (Ecad^+^, detected by immunofluorescence) at 3 hpa. (B) Early nuclear and cytoplasmic *fn1b* expression detected by HCR *in situ* hybridization in wounding epidermis (Ecad^+^, detected by immunofluorescence) at 6 hpa. (C) Late cytoplasmic *fn1b* expression detected by HCR *in situ* hybridization in wounding epidermis (Ecad^+^, detected by immunofluorescence) at 1 dpa. (D) Late cytoplasmic *fn1b* expression detected by HCR *in situ* hybridization in restricted domain within wounding epidermis (Ecad^+^, detected by immunofluorescence) at 2 dpa. (E) Representative H3P^+^ Ecad^+^ *fn1b*^+^ cells for cytometry analysis. (F) Representative H3P^+^ Ecad^+^ *fn1b*^-^ cells for cytometry analysis. (G) Time course analysis of percentage *fn1b*^+^ cells in H3P^+^ Ecad^+^ population, * *FDR<0.05*, Wilcoxon rank sum test. (H) Hole punch assay shows *fn1b* (HCR), Ecad (Ab), and H3P (Ab) stains at 1 dpa in proximal and distal samples at the top. Bottom shows stab injury on the left and uncut control on the right.

We next sought to define the distribution of *fn1b*^+^ basal epidermis on the orthogonal plane, we observed that throughout the time course all *fn1b*^+^ cells always localize to the wounding epidermis where presumably basement membrane was lost after amputation (Supplementary Figure S5A, S5B). Also, there are more *fn1b*^+^ cells in the inter-ray domain compared to the corresponding orthogonal plane along the fin ray, consistent with previous observations in zebrafish^23^ (Supplementary Figure S5A-F). Furthermore, we observed a shift from an earlier time point when *fn1b*^+^ cells represent a multi-layer cell compartment to a single layer compartment over time that delineates the growing tip of the mesenchyme (Figure 5D). We interrogated the proliferative nature of the regeneration-activated *fn1b*^+^ basal epidermis with H3P staining, and observed that few *fn1b*^+^ are positive for H3P. We segmented H3P^+^ cells and we classified them into Ecad^+^ *fn1b*^+^, Ecad^+^ *fn1b*^-^, Ecad^-^ *fn1b*^+^ or Ecad^-^ *fn1b*^-^ cells using cytometry analysis (Figures 5E-F, Supplementary Figure S4C). We quantified the percentage of *fn1b*^+^ cells within the H3P^+^ Ecad^+^ population, and we found that proximal injuries maintain *fn1b* expression within the proliferating epidermis for longer time compared to distal injuries (Figure 5G). This is not the case if we look at the Ecad^-^ compartment (mesenchyme) where we see that the *fn1b*^+^ dividing cells quickly plummet over time independent of amputation position (Supplementary Figure S4D). We measured the spatial distribution of the *fn1b*^+^ Ecad^+^ H3P^+^ cells and found that distal injuries maintain the same spatial distribution across 1 and 2 dpa (Supplementary Figure S4E). However, proximal injuries display a narrower distribution for these cells at 2 dpa (Supplementary Figure S4F). Lastly, we investigated the possibility of interaction between adjacent bones following injury. We performed punch amputation using a 1.5 mm diameter biopsy punch at proximal or distal positions and analyzed *fn1b* expression and H3P at 1 dpa. We observed robust *fn1b* expression all around the circumference of the punch in both proximal and distal injuries with much higher expression at the interface between the wound epidermis and the bone fractures. However, *fn1b* expression was solely localized to fractured bones, which also displayed higher proliferation levels in contrast with adjacent bones that were not amputated (Figure 5H). Interestingly, we observed *fn1b* expression in a stab injury in the absence of exposed mesenchyme, but the expression domain was smaller and weaker compared to punch amputated fins (Figure 5H). Altogether we observe that *fn1b*^+^ basal epidermal cells represent a TRACS with the potential to transduce positional information to the regenerating blastema. Once a blastema is formed (2 dpa), the basal epidermis TRACS loses *fn1b* expression, returning to a homeostatic basal epidermis cell state.

## Discussion

### Amputation position controls at least three regeneration components

The position of amputation along the proximo/distal (P/D) axis of a limb has been shown to influence the response magnitude and amount of pre-existing tissue that participate in the regenerative response. For instance, amputations at different P/D positions lead to differences in growth rate, number of proliferating cells or number of cells expressing a regeneration-induced gene^7,10,24^. It is well-documented that gene expression domains expand in relation to the amputation position, which in turn has been associated with the magnitude of gene expression: the more cells mounting a regeneration response, the larger the expression domain^7,11^. Careful quantitative analyses, however, have provided an alternative interpretation where independent of the number of cells that participate during regeneration, amputation position influences the spatial distribution of these cells. It was shown in axolotls that amputation position determines how far away into the pre-existing tissue progenitor cells will migrate to contribute to the regenerating outgrowth. Supporting the idea that positional information defines the distance from which cells contribute to regeneration^25^.

Our results are consistent with position of amputation affecting both the magnitude and the extent of tissue recruited for regeneration. We found that proliferation profiles are defined by amputation position (Figure 2G). The location of the highest mitotic density (peak proliferation) is proportional to the amputation position, and the size of blastema at 1 dpa also scales linearly with amputation position (Figure 2G-I). We believe both magnitude (*i.e.*, number of dividing cells or number of cells expressing *fgf20*), and location (*i.e.*, peak proliferation at 150 μm from the amputation or cell migration to the blastema) represent two dimensions of regeneration that are influenced by positional information.

Duration of regeneration processes, on the other hand, has been more challenging to address because of the temporal resolution needed to capture regeneration transitions. It has been shown that osteoblast differentiation is accelerated during regeneration^4^, and that cell cycle length can also be altered during regeneration^16,26^. In our study, we captured temporal shifts in regenerative outgrowth acceleration (Figure 1G), tissue-wide proliferation shut-down (Figure 2C), and gene expression (Figure 5G). We observed that these processes last different amounts of time relative to amputation position. The current model considers magnitude and location to be the effectors of positional information during regeneration, but here we present duration as a third component that enables regeneration to respond to positional information. To our knowledge it has not been unambiguously shown that amputation position defines not only the magnitude and extent of tissue involved but also the duration of the biological processes launched during regeneration. Whether there are different mechanisms behind each component, or they share a common regulator remains to be experimentally tested.

### Injury context defines epidermal regeneration cell states

A growing body of evidence indicates that transitional cell states arise following amputation to orchestrate the necessary steps for successful regeneration^27,28^. Injury-induced cellular states have been shown to lead to a spatial compartmentalization of cellular behavior both near and far from the place of insult^29,30,31^. For instance, it has been shown in planarians that Erk signaling propagates tissue-wide during regeneration, and that regeneration ability is lost when its propagation is impaired^31^. Furthermore, in mammalian skin, the cycling characteristics of epidermal progenitor clones are differentially regulated in areas surrounding or away from growing hair follicles, suggesting that location can control spatiotemporal control of cell potencies^32^. We found that upon amputation, fin epidermis transitions into a cell state characterized by a transcriptomic signature enriched in DNA synthesis and ribosome biogenesis genes, irrespective of whether the epidermis was adjacent to an amputation or resided in an uncut fin (Figure 3D, 3H). Our data show that amputation triggers a cell state transition in the epidermis in very distant tissues that were never cut. Interestingly, such cell states were defined by different epidermal subclusters in cut and uncut fins, with the cell state shift of the epidermis in a distant uncut organ restricted to cells that were already committed to undergo cell division (*e.g.*, *pcna*^+^ and *ccna*^+^ cells). We propose that regeneration induces a cell state that primes the epidermis to prepare for rapid proliferation, and the presence of a wound within the same organ pushes the tissue to further recruit epidermal cells that do not show a proliferative phenotype. Whether distant transcriptional responses captured in our study contribute to a feedback mechanism to integrate regenerative signals remains to be investigated.

### Basal epidermis deploys cell states that are dependent on amputation position

We detected small transcriptional changes when comparing scRNAseq data from distal and proximal amputations (Fig 3K, 3L). However, when we compared cell type abundance between distal and proximal amputations, we identified large transcriptional changes in one of the two basal epidermal clusters (Figure 4B, 4C). Because the cells shared cell type identity with basal epidermis but differed in gene expression enrichment depending on whether the fins were cut proximally or distally, we concluded that these differences likely corresponded to different cell states of the same cell type. Moreover, we failed to detect this basal epidermal cell state during fin homeostasis (Figure 4D), indicating that the identified cell state exists only during regeneration.

The identified basal epidermal cell state was characterized by significant enrichment of ECM modifiers and components such as fibronectin (*fn1b*), Wnt ligands, and genes associated with embryonic morphogenesis. Genes encoding ECM components have been shown to be upregulated during regenerative outgrowth^33^. Moreover, disruption of basement membrane components is known to reduce regenerative ability^34^, and transient regeneration-activated cell states (TRACS) have been shown to upregulate gene expression of matrix metalloproteinases that is required for whole body regeneration^27^. Finally, we followed the basal epidermal cell state over time, and found it to be transient (Figure 5A-D and Supplementary Figure 5A-B). Interestingly, the *fn1b*^+^ basal epidermal TRACS uncovered in this study may be orthologous to one of two *fn1b*^+^ epidermis populations observed previously in zebrafish, where early *fn1b*^+^ epidermis was transient and later *fn1b*^+^ epidermis contributed to resident epithelial cells^23^. Our data suggest, therefore, a previously unsuspected role for TRACS in vertebrate regeneration.

## Conclusion

We present a model for the role of the *fn1b*^+^ basal epidermal transient regeneration-activated cell states (TRACS) on reading and writing positional information (Figure 6A-C). In this model, basal epidermis cells adjacent to the amputation plane transition to a TRACS following loss of cell polarity by the absence of basement membrane, and mechanical distension caused by wound closure^12^. We propose that the degree of force applied to the epidermis is proportional to the thickness of the bone at the plane of amputation, along with the corresponding surface area between exposed mesenchyme and wounding epidermis, lead to a proportional number of basal epidermal cells to acquire a basal epidermis TRACS. The transient cell state in turn remodels the extracellular space at the plane of amputation to recruit a proportional number of migrating mesenchyme, resulting in the formation of a concomitantly scaled regeneration blastema. We propose, therefore, that the differences in duration of the proliferative state in proximal versus distal amputations (Figure 2C), the spatial distribution of dividing cell in the tissue (Figure 2G), and the resulting number of the *fn1b*^+^ basal epidermal TRACS along the P/D axis all play a role in codifying the corresponding positional information that results in properly scaled regeneration growth rates. Altogether, our findings have revealed a role for position of amputation in regulating the duration of a proliferative state in injured tissue as well as transient cell states that may relay positional information in animal regeneration.

**Figure 6.**
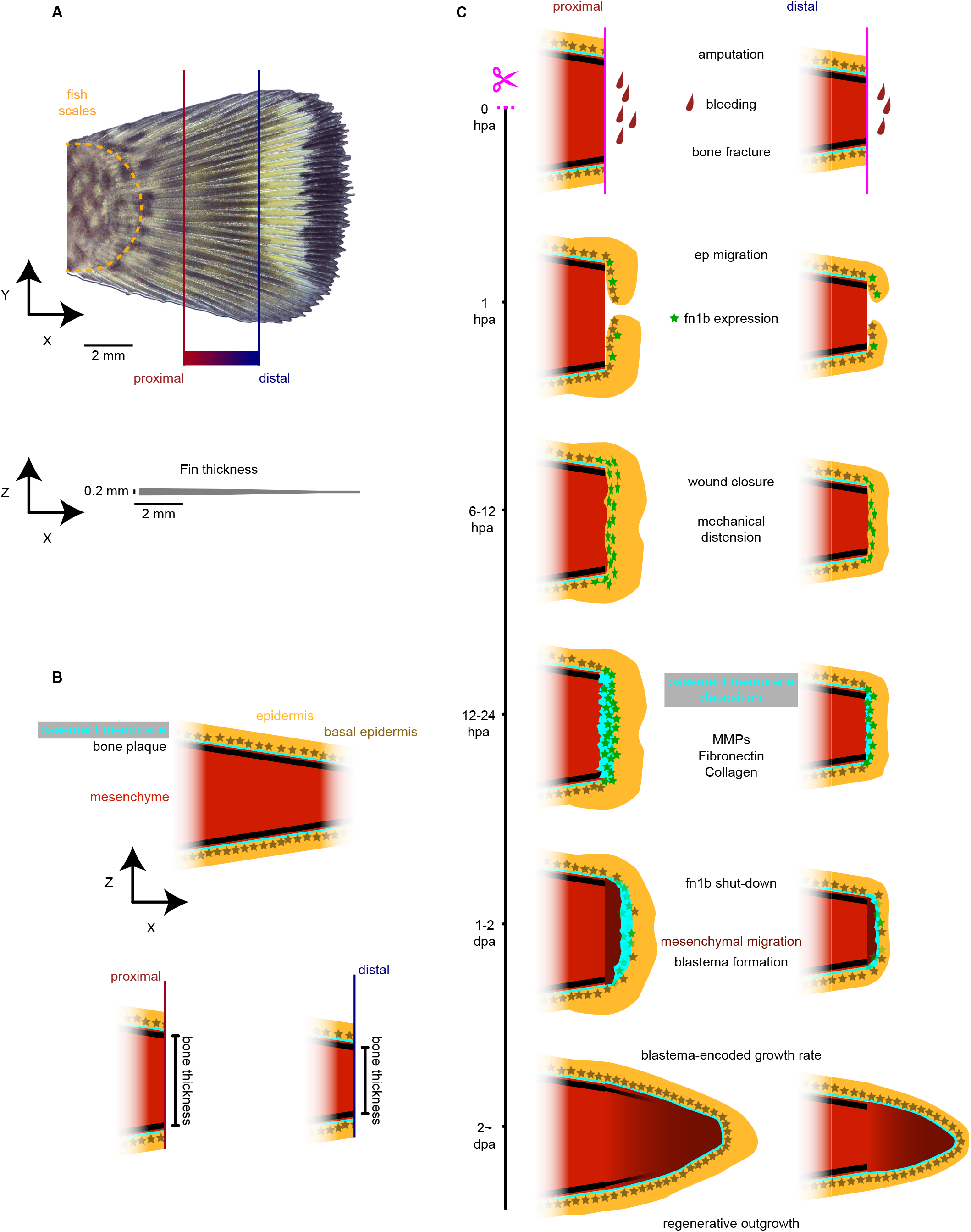
Model for positional information relay during regeneration by basal epidermis. (A) Overview of the fin and proximal and distal definition in this study relative to pigment pattern. Thickness of the fin drawn to scale is showed at the bottom of the image. (B) Color definition for the model presented in (C) and bone thickness comparison between proximal and distal injuries. (C) Model for *de novo* deposition of positional information during regeneration by basal epidermis transient regeneration-activated cell state.

### Limitations of the study

Our study uncovered new dynamics of proliferative activities in injured tissues and the generation of a well-define cell state of basement epeidermis cells in response to injury. However, this reproducible and measurable responses await mechanistic analyses to determine whether or not they are causal to or corrlated with regneration. Still needed are genetic gain or loss of function sttudies. Despite our efforts to genetically manipulate the TRACS, we failed to generate mutants of genes expressed in TRACS that were not embryonic lethal, suggesting the need for conditional mutant alleles for the study of regeneration in adults. Additionally, Our study was solely conducted in males; therefore, further experimentation is needed to test any sex possible differences.

## Acknowledgments

We thank T. Piotrowski, R. Barajas Azpeleta, C. Barradas Chacón, A. Accorsi, V. Doddihal, F. G. Mann, A. Karabulut for critical reading of the manuscript; current and past members of the Sánchez Alvarado lab at SIMR; T. Piotrowski, R. Krumlauf, and N. Rohner for insightful scientific discussion; current and past members of the Technology Centers at SIMR who provided technical support. Vector graphics from Ibrandify (Freepik). We thank the KUMC Genomics Core for generating data on the Illumina NovaSeq 6000 System. The KUMC Genomics Core is supported by the Kansas Intellectual and Developmental Disabilities Research Center (NIH U54 HD 090216), the Molecular Regulation of Cell Development and Differentiation – COBRE (P30 GM122731-03) and the NIH S10 High-End Instrumentation Grant (NIH S10OD021743). This project was funded by the Stowers Institute for Medical Research and the Howard Hughes Medical Institute. Original data underlying this manuscript can be accessed from the Stowers Original Data Repository at http://www.stowers.org/research/publications/libpb-2453

## Author contributions CRediT taxonomy

**Conceptualization**, A.O.G., W.W., A.S.A.; **Methodology**, A.O.G., D.Z., R.R.S., A.R.S., J.R., K.F., J.A.M., B.Y.R.; **Software**, A.O.G., J.R., C.E.B., E.J.R., D.A.A., B.Y.R.; **Validation**, A.O.G.; **Formal Analysis**, A.O.G.; **Investigation**, A.O.G., D.Z., R.R.S., A.R.S., J.R., D.A.A., K.F., B.Y.R., W.W.; **Resources**, D.Z., R.R.S., C.E.B., E.J.R., D.A.A., J.A.M., A.G.P., A.S.A.; **Data Curation**, A.O.G., E.J.R.; **Writing – Original Draft**, A.O.G., A.S.A.; **Writing – Review & Editing**, A.O.G., A.S.A.; **Visualization**, A.O.G.; **Supervision**, A.O.G., R.R.S., A.G.P., W.W., A.S.A.; **Project Administration**, A.O.G., R.R.S., A.G.P., A.S.A.; **Funding Acquisition**, A.G.P., A.S.A.

## Declaration of Interests

The authors declare no competing interests.

## Inclusion and diversity

One or more of the authors of this paper self-identifies as an underrepresented ethnic minority in science. One or more of the authors of this paper self-identifies as a gender minority in science. One or more of the authors of this paper self-identifies as a member of the LGBTQIA+ community. One or more of the authors of this paper received support from a program designed to increase minority representation in science.

## STAR Methods

### RESOURCE AVAILABILITY

#### Lead contact

Requests for reagents and resources should be directed to and will be fulfilled by the lead contact, Alejandro Sánchez Alvarado.

#### Materials availability

This study did not generate new unique reagents, however reagents presented in this study are available from the lead contact with a completed materials transfer agreement and with reasonable compensation by requestor for its processing and shipping.

#### Data and code availability

- Original data underlying this manuscript can be accessed from the Stowers Original Data Repository at http://www.stowers.org/research/publications/libpb-2453. Single cell RNA-seq data have been deposited at GEO and are publicly available under accession number GSE260629 at https://www.ncbi.nlm.nih.gov/geo/query/acc.cgi?acc=GSE260629
- All original code has been deposited at Zenodo and is publicly available under https://doi.org/10.5281/zenodo.10725332
- Any additional information required to reanalyze the data reported in this work is available from the lead contact, Alejandro Sánchez Alvarado.

### EXPERIMENTAL MODEL AND STUDY PARTICIPANT DETAILS

#### Animals

African killifish *Nothobranchius furzeri* were reared at the Stowers Institute and all animal procedures were performed with IACUC approval (Protocol ID: 2022-137). The aquatic animal program meets all federal regulations and has been fully accredited by AAALAC International since 2005. Killifish were single housed in polycarbonate tanks (1.4 liter), with a 14:10 hours light:dark photoperiod. All experiments were performed with the ZMZ1002 wild type line. Only males 2 to 3 months-old were used for our study because of pigment pattern and body size.

### METHOD DETAILS

#### Fin amputation

Experimental fish were anesthetized using 250mg/mL MS-222 (Sigma-Aldrich, Cat. E10521) for 3 mins and placed on top of a plastic petri dish lid (VWR, Cat. 25384-302). Fins were amputated using a disposable razor blade (VWR, Cat.55411-050) or a rapid core punch (World Precision Instruments, Cat. 504647) at the experimental plane of amputation (Supplementary Figure S1A). After amputation the fish was placed back in its housing tank and observed for recovery within 5 minutes.

#### Longitudinal analysis of growth rate, imaging, and quantification

Experimental fish were anesthetized using 250mg/mL MS-222 for 3 mins and placed on their right side on top of a plastic petri dish lid. Glass coverslip (Epredia, Cat. 152222) was placed on top of the tail to avoid water reflection and room temperature system water was added to fully immerse the tail in water. Images were collected every 12 hours for the first 5 days of regeneration and every 24 hours for the remaining of the experiment. Images were acquired with a INFINITY3-6URC (Lumenera) camera on a Leica M205 FCA microscope with a Leica PlanApo 0.63X objective controlled by MicroManager (RRID: SCR_016865)^35,36^, white was balanced with a clean kimwipe (Kimberly Clark, Cat. 34155) and the background was set dark with reflective light coming from both the bottom and the top. After image acquisition the fish was placed back in its housing tank and observed for recovery within 5 minutes.

For image quantification we first integrated the image sequence into a stack that was registered using Linear Stack Alignment with SIFT^37^. The bones to quantify and the base of the fin where the scales end was outlined by hand with the "Straight" and the "Oval" tools in Fiji (RRID: SCR_002285)^38^. The edge of the fin was outlined by traditional thresholding of the images to create a mask, the mask was then subtracted from a one pixel dilate of itself to create a new image with the Edge location as a separate channel on the stack. For each bone the intersection between the oval (fish scales) and the straight ROIs was centered and the image was rotated to align the bone with the x axis. A crop of the preexisting bone tissue was used to register the movie using StackReg^39^ and the registration matrix was propagated to the rest of the stack. The distance between the scales and the bone fracture is defined as the "bone length" and the distance between the bone fracture and the edge of the tissue is defined as the "regeneration length". Growth rate measurements were done by subtracting the average regeneration length on one timepoint from the antecedent timepoint and dividing the result by the length of time between the two timepoints, growth rate is expressed in millimeters per day. Changes in growth rate over time is defined as "growth acceleration", and it was calculated using a linear regression of at least three timepoints to measure the slope of the linear model.

#### Sample collection and bleaching for IF, HCR or HCR-IF

Fish were euthanized for 5 mins with 500mg/L MS-222 followed by hypothermic shock in 4°C cold system water for 30 mins, in between the MS-222 treatment and the hypothermic shock, fin of interest was collected and fixed with 4% PFA (Electron Microscopy Sciences, Cat. 15710) 0.1% Tween-20 1X PBS for 4 hours at room temperature. The fins were dehydrated in 25%-50%- 75%-100% EtOH (Sigma-Aldrich, Cat. E7023) train in PBST (0.1% Tween-20 1X PBS) for at least 4 hours per step. Fins were left for at least one night in 100% EtOH (200 proof), this step is very important for enabling reagent penetration and successful bleaching. A first bleaching step was done with 6% H2O2 (Sigma-Aldrich, Cat. H1009) 80% EtOH under an LED lamp (3W) for twelve hours at room temperature, then rehydration was done on the reverse train 50%-25%-EtOH- PBST for at least 4 hours per step. A second bleaching step was done with 3% H_2_O_2_ 5% formamide (Sigma-Aldrich, Cat. S4117) 0.1% Tween-20 in 1X SSC under an LED lamp (3W) for 1 hour at room temperature. Two 5-min washes were done with PBST before proceeding with staining.

#### IF staining

Following bleaching, samples were blocked overnight with 10% Goat Serum (Thermo Fisher, Cat. 16210072), 5% Western Blocking Reagent solution (Sigma-Aldrich, Cat. 11921673001), 2.5% Horse Serum (Thermo Fisher, Cat. 26050070), 5% DMSO (Sigma-Aldrich, Cat. 472301), 123 mM NaN_3_ (Sigma-Aldrich, Cat. S2002) in PBST. Two 5-min washes with PBST were done after blocking and the samples were incubated with primary antibody solution: anti-H3P at 1:400 dilution (Cell Signaling Technology, catalog number: 3377S, RRID: AB_1549592), anti-Ecad at 1:200 dilution (BD Biosciences, catalog number: 610182, RRID: AB_397581) diluted in 10% FBS (Thermo Fisher, Cat. A5256801), 5% DMSO, 123 mM NaN_3_ in PBST for 24 hours at room temperature in roller shaker (Cole-Parmer, Cat. UX-51901-23). Four 20-minutes washes with PBST at room temperature were done in between primary and secondary antibody incubation. Secondary antibody solution: anti-rabbit AF488 at 1:400 dilution (Thermo Fisher, Cat. A48286), anti-mouse AF555 at 1:500 dilution (Thermo Fisher, Cat. A48287) diluted in 10% FBS, 5% DMSO, 123 mM NaN_3_ in PBST for 24 hours at room temperature in roller shaker protected from ambient light. Six 20-minutes washes with PBST at room temperature were done afterwards. Nuclear staining was done with 1μg/mL DAPI (Thermo Fisher, Cat. 62248), or 1nM YOYO-1 Iodide (Thermo Fisher, Cat. Y3601) in PBST overnight at room temperature in roller shaker, and two 20-minute washes with PBST at room temperature were done prior to clearing. Overnight tissue clearing was done with 1mL of EasyIndex (LifeCanvas Technologies) and the sample was mounted on round coverslip bottom petri dishes (Fisher Scientific, Cat. 50-305-805) in the same medium that was used for clearing, an additional glass coverslip was placed on top of the sample and volume was minimized to get the sample to lay flat at the bottom of the petri dish.

#### HCRv3-IF staining

Following bleaching, samples were pre-hybridized with 500μL probe hybridization buffer (Molecular Instruments) preheated at 37°C for 30 min. In the meantime, probe solution was prepared by adding 4μL of each probe set to 500 μL hybridization buffer preheated at 37°C. Hybridization was done for 24 hours at 37°C in rocking shaker (Fisher Scientific, Cat. 88-861-025) with the tubes oriented vertically on a rack so the tissue is not damaged. Samples were washed 4 times for 15 minutes with probe wash buffer (Molecular Instruments) at 37°C in rocking shaker, followed by 2 washes for 15 minutes each with SSCT at room temperature in roller shaker. Pre-amplification was done with 500 μL of room temperature amplification buffer (Molecular Instruments) for 30 mins in roller shaker. In the meantime, snap cooling of h1 and h2 harpins corresponding to AF647 and AF546, were incubated at 95°C for 90 secs and then cooled at room temperature for 30 mins in the dark. Amplification solution was prepared by adding h1 and h2 harpins in 500 μL amplification buffer at room temperature. Samples were incubated in amplification solution for 24 hours at room temperature in roller shaker, washed 5 times with 0.01% Tween-20 1X SSC, and washed once with PBST for 15 minutes per wash at room temperature in roller shaker. Samples were fixed with 4% PFA 0.1% tween PBST for 1 hour at room temperature and washed once with PBST for 15 minutes at room temperature in roller shaker. For HCR-only samples we counterstained with nuclear stain, cleared and mounted the same way than IF (see above). For immunofluorescence following HCR we skipped the blocking step and went straight to primary antibody incubation and followed the IF staining protocol (see above).

#### Confocal imaging

Images were acquired with an Orca Flash 4.0 sCMOS 100fps at full resolution on a Nikon Eclipse Ti microscope equipped with a Yokagawa CSU W1 10,000 rpm Spinning Disk Confocal with 50 μm pinholes. Samples were illuminated with 405nm(5.73mw), 488nm(6.75mw), 561nm(5.84mw) and 640nm(6.62mw) lasers (LUNV 6-line Laser Launch) with nominal power measures at the objective focal plane. This spinning disk confocal is equipped with a quad dichroic filter for excitation with 405/488/561/640nm. Emissions filters used to acquire this image were 430-480 nm for DAPI or YOYO-1, 507-543 nm for AF488, 570-640 nm for AF555 or AF546 and 662.5-737.5 nm for AF647.

For figure 2, we used a Nikon Plan Apochromat Lambda 10x objective lens, N.A. 0.45, 0.78 μm/px objective with 200 ms exposure for IF channels, and 100 ms for DAPI channel at 3 μm z resolution. For figures 4, 5, supplementary figures S2, S4 and S5 high magnification images we used a Nikon Plan Apochromat Lambda LWD 40x objective lens, N.A 1.15, 0.283 μm/px with 300 ms exposure for HCR channels and 100 ms for DAPI channel at 3 μm z resolution. For figure 5 low magnification images we used a Nikon Plan Apochromat Lambda 4x objective lens, N.A. 0.2, 1.735 μm/px with 200 ms exposure for HCR channels and 50 ms for YOYO-1 channel at 25 μm z resolution.

#### H3P segmentation and cytometry analysis

Tiled images were stitched using the Grid/Collection stitching Plugin^40^ from Fiji, and X, Y, and Z coordinates were identified for H3P positive nuclei segmented following the pipeline published elsewhere^41^. Quadrants in supplementary figure S4C (cytometry analysis) was done using fluorescence minus one (FMO) negative *fn1b* (HCR) and Ecad (IF) controls.

#### Proliferation profile build

The analysis starts with segmentation of the tail into regions—each region contains exactly one ray and number of regions is equal to number of rays. The ray is determined by slope and intercept of the line determining the bone in the Cartesian coordinates. The i-th region boundaries (for all regions except the first and the last) are computed as a straight line representing a bisector cutting the angle between the i-th and (i+1)-th rays. The upper boundary of the first (upper) region and the lower boundary of the last (lowest) region are computed to make the rays inside the region to be its bisector. For each region we find all cells with coordinates X, Y belonging to this part of the plane using the data from the segmentation step (see above).

As region boundaries are not parallel lines, we can determine their intersection point that is served as the region center. Using this point we convert all X, Y coordinates of each point of interest (cells, scales, edge points, amputation points etc.) into two other numbers r and ‘Y to bone’ (the last one is ready for the output). The radial coordinate r is determined as a distance between the center and the given point, while the ‘Y to bone’ coordinate is measured as distance from the point to the bone along the normal to the bone. We use the ray line to compute the distance between two points along this line.

Note that there are two types of regions – the central regions that have amputation points and the side regions free of such points. The regions of both types are characterized by the sequence of edge points. These special points are used for a special (scaled) measure of the cell position. Specifically, when the region has amputation points, we find the average radial distance between the region center and all amputation points. When we use the edge points, we generate a polynomial approximation R(⍰) of the curve corresponding to the ordered sequence of the edge points in the region boundary. The function R(⍰) determines the dependence of the edge point radial distance from the region center as a function of the angular polar coordinate ⍰.

Then the following distances along the ray are computed: from the cell to the edge, from the cell to the averaged amputation position, these values allow to produce ‘r bone Amp’ and ‘r bone Edge’. We also find the distance from the scale center to the edge, from the bifurcation point to the edge, from the amputation point to the edge to be added to corresponding output files.

Once we generated distance measurements of each given H3P positive nuclei to the amputation plane parallel to the bone, we bin the bone axis every 50 μm, and we counted the H3P positive nuclei within each bin to generate heatmaps of proliferation distribution (Figure 2C). Next, we iteratively slide the 50 μm bin for 1 μm at a time along the bone axis to generate a proliferation density profile along the bone dimension, the peak proliferation (Figure 2G), corresponds to the highest value of the proliferation density profile after a median blur (kernel=3).

#### Single cell dissociation, CellPlex labeling and flow cytometry

Fish were euthanized at the expressed time point and tissues of interest were rinsed in cold 0.1% BSA (Fisher Scientific Cat. BP9706100) 1X PBS (PBS-BSA). The tissue was minced using a fine blade and single cell suspension was achieved by dissociation in 10mL 1mg/mL collagenase type 2 (Worthington Biochemical, Cat. LS004174) for 5mins at 37dC followed by 70 μm filtration (Fisherbrand, Cat. 22-363-548). The cells were rinsed by adding 25mL 0.1% BSA 1X PBS (PBS-BSA) and centrifuged for 3 minutes at 500g at 4C with slow deacceleration (slow deacceleration will help keep the pellet intact).

A small aliquot of the cells was counted using 500 nM Draq5 (Biostatus, Cat. DR50200) solution in an EC800 flow cytometer. For regular 10x run, cells were stained with 1μg/mL DAPI, 500 nM Draq5 in PBS-BSA at 4e6 cells/mL and sorted as described below. For CellPlex 10x run, cells were labeled with 3⍰ CellPlex Kit Set A (10x Genomics, PN-1000261) according to the manufacturer’s directions. A small aliquot was counted one more time using 500 nM Draq5 solution in an EC800 flow cytometer, and different samples were pooled together to balance the cell pool equally across samples. Then cells were stained with DAPI-Draq5 (same as above) at 4e6 cells/mL and sorted as described below. We employed CellPlex to collect biological replicates of all our conditions as well as prevent batch effects between samples within a multiplex experiment.

For cell sorting, the sample buffer and the cell sorter sheath fluid contained 0.1% Poloxamer 188 (Sigma-Aldrich, Cat. P5556) to reduce shear stress and improve post-sort cell viability. Sorting of live cells was performed on a 6-laser BD S6 FACSymphony equipped with a 100-μm nozzle, and chilled to 4°C at all times. Because the Draq5-DAPI staining produced no spillover into the target detectors on this cell sorter, no compensation was performed. Due to the heterogeneous nature of the sample, an FSC/SSC scatter gate was not used to avoid biasing the sorted sample for specific cell types, instead we triggered acquisition with Draq5 to capture all nucleated events. The Draq5-DAPI staining pattern displayed minor shifts over the course of the sort, which was accounted for by adjustments of the sort gate. To ascertain sample quality, a post-sort viability assay was performed on the BD S6 FACSymphony prior to further processing of each sample. The sorter was flushed with sample buffer (cold 0.1% BSA 1X PBS) for 1 minute between samples to remove leftover nuclear stain and prevent secondary staining. Samples with post-sort viability >96% were used for library preparation and sequencing.

#### scRNAseq library preparation and sequencing

##### For regular 10X

One proximal and one distal single cell library were generated using conventional methods. Dissociated, sorted cells were assessed for concentration and viability via Luna-FL cell counter (Logos Biosystems). Cells deemed to be at least 97% viable were loaded on a Chromium Single Cell Controller (10x Genomics), based on live cell concentration. Libraries were prepared using the Chromium Next GEM Single Cell 3’ Reagent Kits v3.1 (10x Genomics) according to manufacturer’s directions. Resulting cDNA and short fragment libraries were checked for quality and quantity using a 2100 Bioanalyzer (Agilent Technologies) and Qubit Fluorometer (Thermo Fisher Scientific). With cells captured estimated at ∼8,000-9,000 cells per sample, libraries were pooled and sequenced to a depth necessary to achieve at least 42,000 mean reads per cell on an Illumina NovaSeq 6000 instrument utilizing RTA and instrument software versions current at the time of processing with the following paired read lengths: 28*10*10*90bp.

##### For CellPlex 10x

Three multiplexed libraries were generated encompassing: 1) regeneration and homeostasis samples, 2) dorsal samples, and 3) proximal and distal samples. The CellPlex labeled, pooled cell samples were assessed for concentration and viability via Luna-FL cell counter (Logos Biosystems). Samples with cells deemed to be at least 97% viable were loaded on a Chromium Single Cell Controller (10x Genomics), based on live cell concentration targeting 20,000, 30,000 or 50,000 cells. Libraries were prepared using the Chromium Next GEM Single Cell 3’ Reagent Kits v3.1 with Feature Barcode technology for Cell Multiplexing (10x Genomics) according to manufacturer’s directions. Resulting cDNA, short fragment libraries, and CMO libraries were checked for quality and quantity using a 2100 Bioanalyzer (Agilent Technologies) and Qubit Fluorometer (Thermo Fisher Scientific). Multiplexed gene expression and CMO libraries were pooled, as specified by manufacturer, and sequenced to a depth necessary to achieve at least 26,000 mean reads per cell on an Illumina NovaSeq 6000 instrument utilizing RTA and instrument software versions current at the time of processing with the following paired read lengths: 28*10*10*90bp

#### scRNAseq alignment primary analysis

Fastqs were demultiplexed using cellranger mkfastq (10x Genomics Cell Ranger^42^ 6.0.1) with default settings. Cellranger multi (10x Genomics Cell Ranger 6.0.1) was used to align the fastqs against Ensembl^43^ 104 Nfu_20140520 and filter, count barcodes and UMIs. Feature-barcode matrices were analyzed as described below.

#### scRNAseq quality control, integration, differential expression analysis and GO enrichment

Demultiplexed sample feature count matrices were loaded into R using Seurat^44^ (Seurat 4.3.0), mitochondrial percentages were calculated using the PercentageFeatureSet function and a *median + 2 sd* threshold was use to filter out cells with high mitochondrial gene expression, the resulting percent.mt distribution in the integrated object had a median of 3.9% and the highest value was 21.5%. Low nFeature_RNA count was filtered with *median - 1.2 sd* and high nFeature_RNA count was filtered with *median + 4 sd*, the resulting nFeature_RNA distribution in the integrated object had a median of 1536 and the highest value was 5259 nFeature_RNA. We decided to apply relative statistical thresholds to compensate between batch effects and sequencing depth between samples considering doublets were removed during demultiplexing on the previous step. After quality control, all samples were normalized individually using the SCTransform function regressing percent.mt from the model. All the samples for integration were put on a list and anchor features were identified with SelectIntegrationFeatures (nfeatures = 8000), samples were prepared for integration using the function PrepSCTIntegration and the anchor features, and integration anchors were identified with the function FindIntegrationAnchors using the list containing the samples, SCT as the normalization method and the anchor features. Finally, all samples were integrated using the function IntegrateData with the integration anchors and SCT as the normalization method. Once integrated, RunPCA, RunUMAP and FindNeighbors functions were run on the *integrated* assay, and clusters were computed with the FindClusters function by iteratively changing the cluster resolution from 0.2 to 2 in 0.1 intervals. Cell markers of differential expression analysis was done using the FindMarkers function and selecting the subset of cells to compare to each other. GO enrichment was performed using the enrichGO function from the clusterProfiler package^45^ (clusterProfiler 4.6.2) and a custom killifish database built with the makeOrgPackage function from the AnnotationForge package^46^. We used a corresponding table that connects Ensembl IDs to their corresponding Gene Ontology terms pulled down from Ensembl’s BioMart tool^47^ (Ensembl 104).

#### Differential abundance analysis using Milo

To understand the enrichment of certain cell types between conditions, differential abundance between the cell neighborhoods for a given condition were tested using miloR (v0.1.0)^17^. We tested for differential abundance between the conditions (proximal vs distal, regenerated vs homeostasis). Using Seurat generated graph, a K-nearest neighborhood (KNN) graph was precomputed which assigned the cells to a neighborhood with the parameters (k=20, d=30, prop = 0.1). Cells within each neighborhood for a given sample were then counted and tested for differential abundance using a generalized linear design framework while accounting for multiple comparison testing using the spatial FDR in miloR. We then annotated the neighborhoods with the cell clusters and neighborhoods with a fraction less than 0.7 were annotated to be “mixed”.

#### Quantification and statistical analysis

We used R^48^ (version 4.2.3) to do statistical analysis. All numbers of individuals used for the experiments and the statistical test method used can be found in the corresponding figure legends. Generally, we assigned the significance level based on p value in following manner: *\* p* < 0.05; ** p < 0.01, *** p < 0.001.

## Supplemental information

**Supplementary Figure S1. Fish size and growth rate comparison.**

(A) Representative image of the male caudal fin in killifish, the boundary between the scales and the caudal fin is delineated by the orange dotted line. The distal amputation is performed between the spotted pigmentation region and the yellow pigmentation line perpendicular to the anterior and posterior axis. The proximal amputation is performed at the coloring transition from bright to dark within the spotted pigmentation region perpendicular to the anterior and posterior axis. Both cuts correspond to the second and first bifurcation of the fin rays respectively.

(B) Back calculation of growth rate measurements in zebrafish published by Uemoto et.al. 2020 superimposed with killifish regeneration growth rate of equivalent size fish.

(C) Tail fin length of individual fish used in Figure 1 measured from the base of the fin to the edge of the tissue prior to amputation.

(D) Anterior-Posterior body length of individual fish used in Figure 1 measured from the mouth to the base of the fin.

(E) Bone length measured from the base of the fin where the scales meet the tail to the amputation plane of all bones that support the model in Figure 1.

ns not significant, ∗∗∗ *p<0.001*, Wilcoxon rank sum test.

**Supplementary Figure S2. Mitotic cells mainly correspond to dividing epidermis.**

(A) 24 hpa bone and inter-ray 10 μm MAX projections of orthogonal views of high-magnification confocal stacks.

(B) Number of H3P^+^ nuclei inside 2 mm window from the amputation plane along the bone axis. (C) Number of H3P^+^ nuclei inside 0.5 mm window from the amputation plane along the bone axis.

(C) Number of H3P^+^ nuclei inside 1 mm window counting from 1 mm from the amputation plane to 2 mm from the amputation plane along the bone axis.

(D) Definition of distance to amputation used to calculate proliferation profiles and peak proliferation.

**Supplementary Figure S3. CellPlex workflow and single cell atlas cell type definition.**

(A) Multiplexed scRNAseq workflow using 10X CellPlex reagents.

(B) Dimensional reduction UMAP plot of the integrated dataset with cell type definitions clustered at resolution 1.0, colors are randomly selected to each cluster.

(C) Heatmap of cell type markers.

(D) Dimensional reduction UMAP distribution of regeneration and homeostasis cells captured within the caudal clusters (CC) and dorsal clusters (DC) in both caudal and dorsal datasets.

(E) *ccna*, *mki67* and *pcna* expression within the integrated dataset.

(F) Heatmap of enrichment analysis between RRG and caudal, dorsal, proximal, and distal cells within caudal clusters (CC) and dorsal clusters (DC). Fisher’s exact test, *FDR < 0.001*.

(G) Heatmap of regeneration upregulated genes within caudal clusters (CC) and dorsal clusters (DC) split by any bias towards one or the other cluster group. and top GO terms associated with the gene subset (*FDR < 0.05*). Cells were randomly sampled to balance all four sample groups.

**Supplementary Figure S4. *fn1b*^+^ expression shuts down at later time points,*fn1b* and Ecad cytometry analysis, and spatial distribution of *fn1b*^+^ Ecad^+^ H3P^+^ cells.**

(A) 3 dpa *fn1b* (HCR), Ecad (Ab), H3P (Ab) whole mount staining. (needs a scale bar) (B) 5 dpa *fn1b* (HCR), Ecad (Ab), H3P (Ab) whole mount staining.

(B) Ecad and *fn1b* cytometry analysis on H3P^+^ cells.

(C) % of *fn1b*^+^ cells within the Ecad^-^ H3P^+^ population over time.

(D) Spatial distribution of *fn1b*^+^ Ecad^+^ H3P^+^ cells in distal injuries at 1 and 2 dpa. (F) Spatial distribution of *fn1b*^+^ Ecad^+^ H3P^+^ cells in proximal injuries at 1 and 2 dpa.

**Supplementary Figure S5. Orthogonal analysis of regeneration time course.**

(A) Ray and inter-ray orthogonal views of regenerating proximal and distal samples at 3 hpa *fn1b* (HCR), Ecad (Ab), H3P (Ab) whole mount staining. (fix scale bar)

(B) Same as in (A) at 6 hpa *fn1b* (HCR), Ecad (Ab), H3P (Ab) whole mount staining.

(C) Same as in (A) at 1 dpa *fn1b* (HCR), Ecad (Ab), H3P (Ab) whole mount staining.

(D) Same as in (A) at 2 dpa *fn1b* (HCR), Ecad (Ab), H3P (Ab) whole mount staining.

(E) Ray orthogonal views of regenerating proximal and distal samples at 3 dpa *fn1b* (HCR), Ecad (Ab), H3P (Ab) whole mount staining.

(F) Same as in (D) at 5 dpa *fn1b* (HCR), Ecad (Ab), H3P (Ab) whole mount staining.

**Supplementary Table. Differential gene expression between 12hpa vs homeostasis, and proximal vs distal within CC and DC.**

Left side of the spreadsheet shows differentially expressed genes and right side of the spreadsheet shows statistically enriched GO terms (*padj<0.05*). All data corresponds to heatmaps in main Figure 3D, 3H, 3K and 3L.

## Notes

### Competing Interest Statement

The authors have declared no competing interest.

