## Supplemental Figures for "Positional information modulates transient regeneration-activated cell states during vertebrate appendage regeneration"

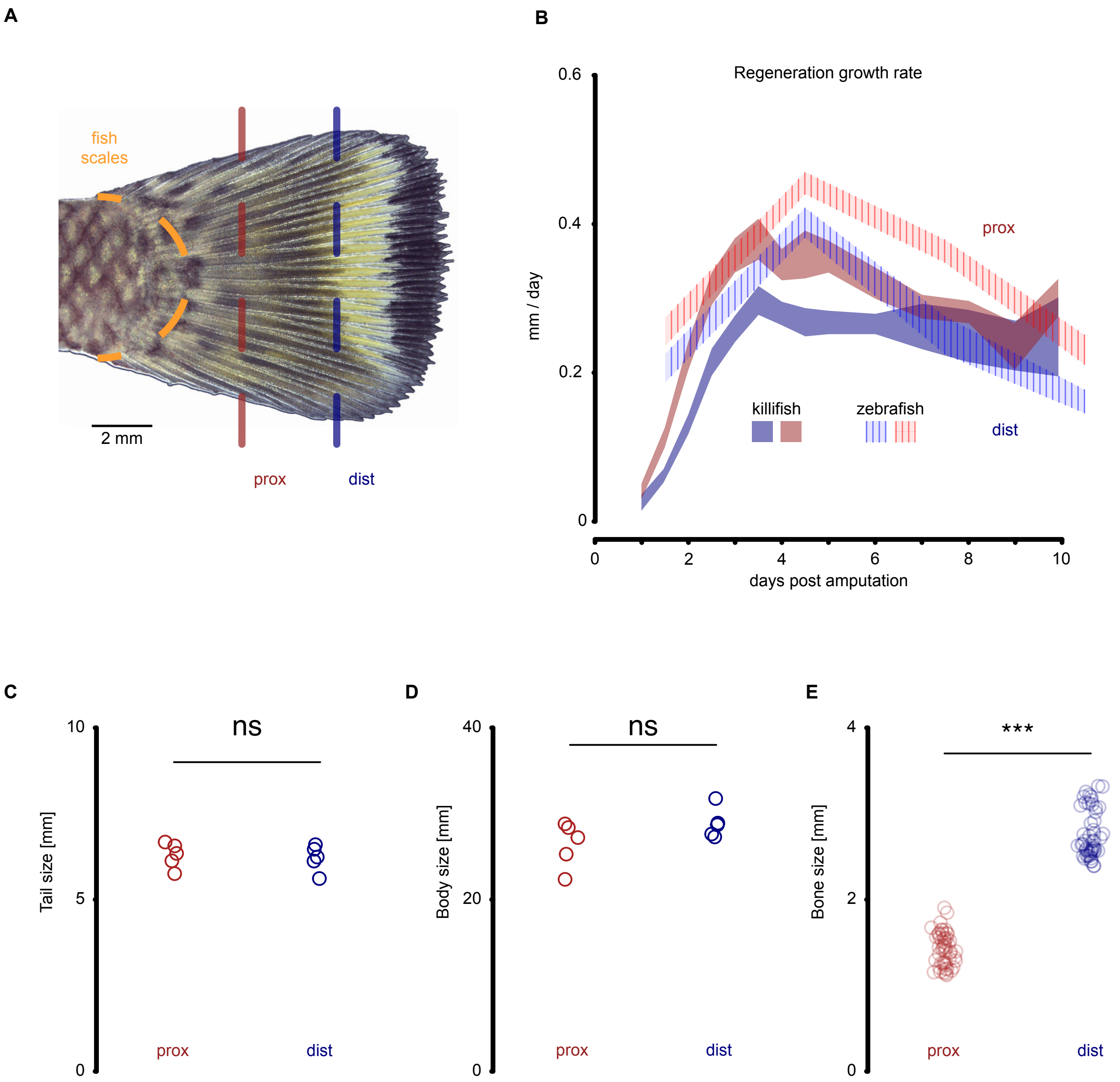
A
fish
scales
2 mm
prox
dist
B
Regeneration growth rate
0.6
0.4
0.2
0
mm / day
prox
killifish
zebrafish
dist
0
2
4
6
8
10
days post amputation
C
10
5
0
Tail size [mm]
ns
prox
dist
D
40
20
0
Body size [mm]
ns
prox
dist
E
4
2
0
Bone size [mm]
***
prox
dist

**Supplementary Figure S1. Fish size and growth rate comparison.**
(A) Representative image of the male caudal fin in killifish, the boundary between the scales and the caudal fin is delineated by the orange dotted line. The distal amputation is performed between the spotted pigmentation region and the yellow pigmentation line perpendicular to the anterior and posterior axis. The proximal amputation is performed at the coloring transition from bright to dark within the spotted pigmentation region perpendicular to the anterior and posterior axis. Both cuts correspond to the second and first bifurcation of the fin rays respectively.
(B) Back calculation of growth rate measurements in zebrafish published by Uemoto et.al. 2020 superimposed with killifish regeneration growth rate of equivalent size fish.
(C) Tail fin length of individual fish used in Figure 1 measured from the base of the fin to the edge of the tissue prior to amputation.
(D) Anterior-Posterior body length of individual fish used in Figure 1 measured from the mouth to the base of the fin.
(E) Bone length measured from the base of the fin where the scales meet the tail to the amputation plane of all bones that support the model in Figure 1.

ns not significant, *** $p<0.001$, Wilcoxon rank sum test.

A

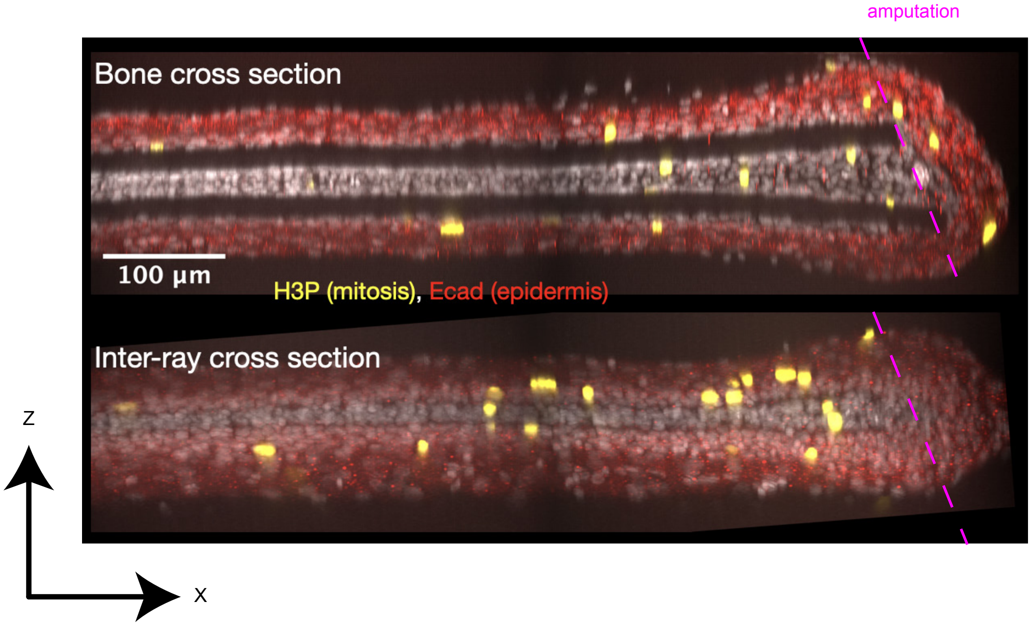

amputation
Bone cross section
100 µm
H3P (mitosis), Ecad (epidermis)
Inter-ray cross section
Z
X


B

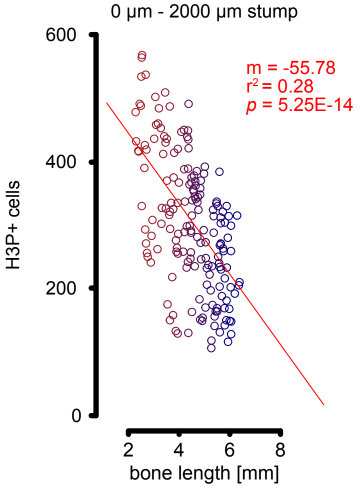

0 µm - 2000 µm stump
m = -55.78
$r^2$ = 0.28
$p$ = 5.25E-14
H3P+ cells
bone length [mm]


C

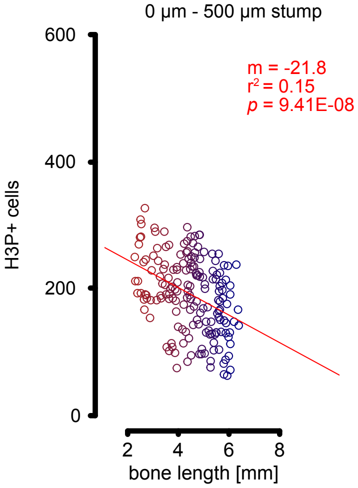

0 µm - 500 µm stump
m = -21.8
$r^2$ = 0.15
$p$ = 9.41E-08
H3P+ cells
bone length [mm]


D

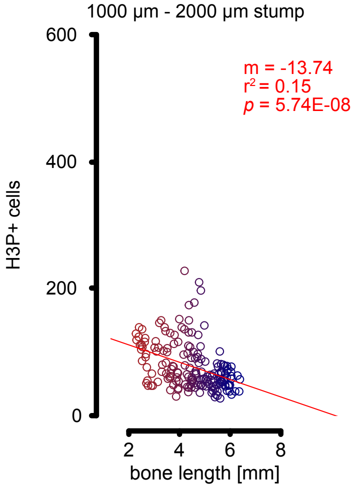

1000 µm - 2000 µm stump
m = -13.74
$r^2$ = 0.15
$p$ = 5.74E-08
H3P+ cells
bone length [mm]


E

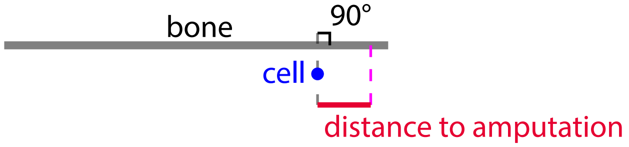

bone
90°
cell
distance to amputation

**Supplementary Figure S2. Mitotic cells mainly correspond to dividing epidermis.**

(A) 24 hpa bone and inter-ray 10 µm MAX projections of orthogonal views of high-magnification confocal stacks.

(B) Number of $H3P^+$ nuclei inside 2 mm window from the amputation plane along the bone axis.

(C) Number of $H3P^+$ nuclei inside 0.5 mm window from the amputation plane along the bone axis.

(D) Number of $H3P^+$ nuclei inside 1 mm window counting from 1 mm from the amputation plane to 2 mm from the amputation plane along the bone axis.

(E) Definition of distance to amputation used to calculate proliferation profiles and peak proliferation.

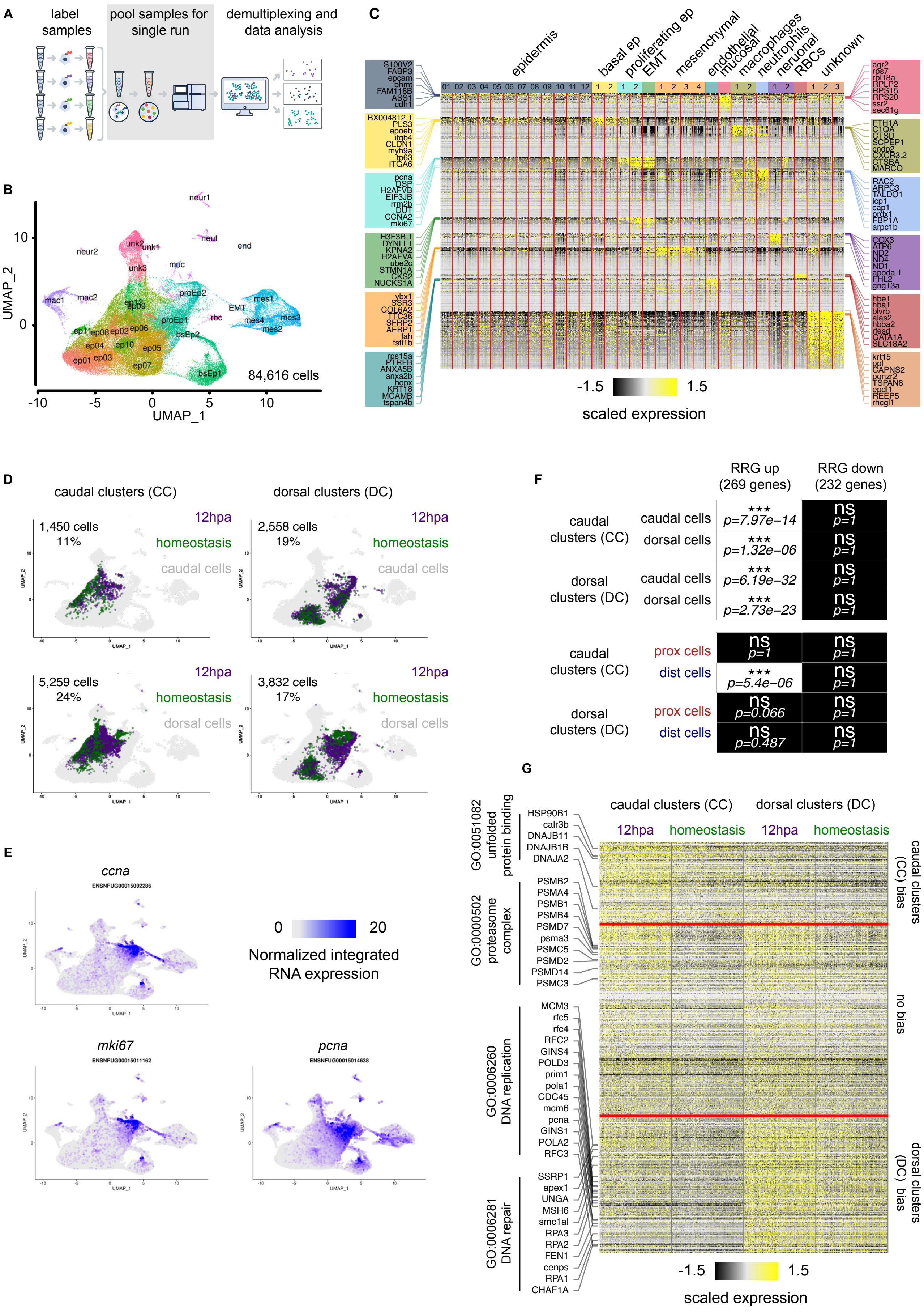
A
label samples
pool samples for single run
demultiplexing and data analysis
B
UMAP_2
UMAP_1
84,616 cells
neur1
neur2
unk2
unk1
neut
end
unk3
muc
mac1
mac2
proEp2
mes1
EMT
rbc
mes3
mes4
mes2
proEp1
bsEp2
bsEp1
ep01
ep02
ep03
ep04
ep05
ep06
ep07
ep08
ep09
ep10
ep11
ep12
C
epidermis
basal ep
proliferating ep
EMT
mesenchymal
endothelial
mucosal
macrophages
neutrophils
neuronal
RBCs
unknown
S100V2
FABP3
epcam
bhmt
FAM118B
ASS1
cdh1
BX004812.1
PLS3
apoeb
itgb4
CLDN1
myh9a
tp63
ITGA6
pcna
DSP
H2AFVB
EIF3JB
rrm2b
DUT
CCNA2
mki67
H3F3B.1
DYNLL1
KPNA2
H2AFVA
ube2c
STMN1A
CKS2
NUCKS1A
vbx1
SSR3
COL6A2
TTC36
SFRP2
AEBP1
fah
fstl1b
rps15a
PTRFB
ANXA5B
anxa2b
hopx
KRT18
MCAMB
tspan4b
agr2
rps7
rpl18a
RPLP2
RPS15
RPS20
ssr2
sec61g
FTH1A
C1QA
CTSD
SCPEP1
cndp2
CXCR3.2
CTSBA
MARCO
RAC2
ARPC3
TALDO1
lcp1
cap1
prdx1
FBP1A
arpc1b
COX3
ATP6
ND2
ND4
ND1
apoda.1
FHL2
gng13a
hbe1
hba1
blvrb
alas2
hbba2
rfesd
GATA1A
SLC18A2
krt15
ppl
CAPNS2
ponzr2
TSPAN8
epdl1
REEP5
rhcgl1
-1.5
1.5
scaled expression
D
caudal clusters (CC)
dorsal clusters (DC)
caudal cells
dorsal cells
1,450 cells
11%
2,558 cells
19%
5,259 cells
24%
3,832 cells
17%
12hpa
homeostasis
caudal cells
dorsal cells
E
ccna
ENSNFUG00015002286
mki67
ENSNFUG00015011162
pcna
ENSNFUG00015014638
0
20
Normalized integrated RNA expression
F
RRG up (269 genes)
RRG down (232 genes)
caudal clusters (CC)
caudal cells
*** p=7.97e−14
ns p=1
dorsal cells
*** p=1.32e−06
ns p=1
dorsal clusters (DC)
caudal cells
*** p=6.19e−32
ns p=1
dorsal cells
*** p=2.73e−23
ns p=1
caudal clusters (CC)
prox cells
ns p=1
ns p=1
dist cells
*** p=5.4e−06
ns p=1
dorsal clusters (DC)
prox cells
ns p=0.066
ns p=1
dist cells
ns p=0.487
ns p=1
G
caudal clusters (CC)
dorsal clusters (DC)
12hpa
homeostasis
GO:0051082 unfolded protein binding
HSP90B1
calr3b
DNAJB11
DNAJB1B
DNAJA2
GO:0000502 proteasome complex
PSMB2
PSMA4
PSMB1
PSMB4
PSMD7
psma3
PSMC5
PSMD2
PSMD14
PSMC3
GO:0006260 DNA replication
MCM3
rfc5
rfc4
RFC2
GINS4
POLD3
prim1
pola1
CDC45
mcm6
pcna
GINS1
POLA2
RFC3
GO:0006281 DNA repair
SSRP1
apex1
UNGA
MSH6
smc1al
RPA3
RPA2
FEN1
cenps
RPA1
CHAF1A
caudal clusters (CC) bias
no bias
dorsal clusters (DC) bias
-1.5
1.5
scaled expression

**Supplementary Figure S3. CellPlex workflow and single cell atlas cell type definition.**
(A) Multiplexed scRNAseq workflow using 10X CellPlex reagents.
(B) Dimensional reduction UMAP plot of the integrated dataset with cell type definitions clustered at resolution 1.0, colors are randomly selected to each cluster.
(C) Heatmap of cell type markers.
(D) Dimensional reduction UMAP distribution of regeneration and homeostasis cells captured within the caudal clusters (CC) and dorsal clusters (DC) in both caudal and dorsal datasets.
(E) *ccna*, *mki67* and *pcna* expression within the integrated dataset.
(F) Heatmap of enrichment analysis between RRG and caudal, dorsal, proximal, and distal cells within caudal clusters (CC) and dorsal clusters (DC). Fisher's exact test, $FDR < 0.001$.
(G) Heatmap of regeneration upregulated genes within caudal clusters (CC) and dorsal clusters (DC) split by any bias towards one or the other cluster group. and top GO terms associated with the gene subset ($FDR < 0.05$). Cells were randomly sampled to balance all four sample groups.

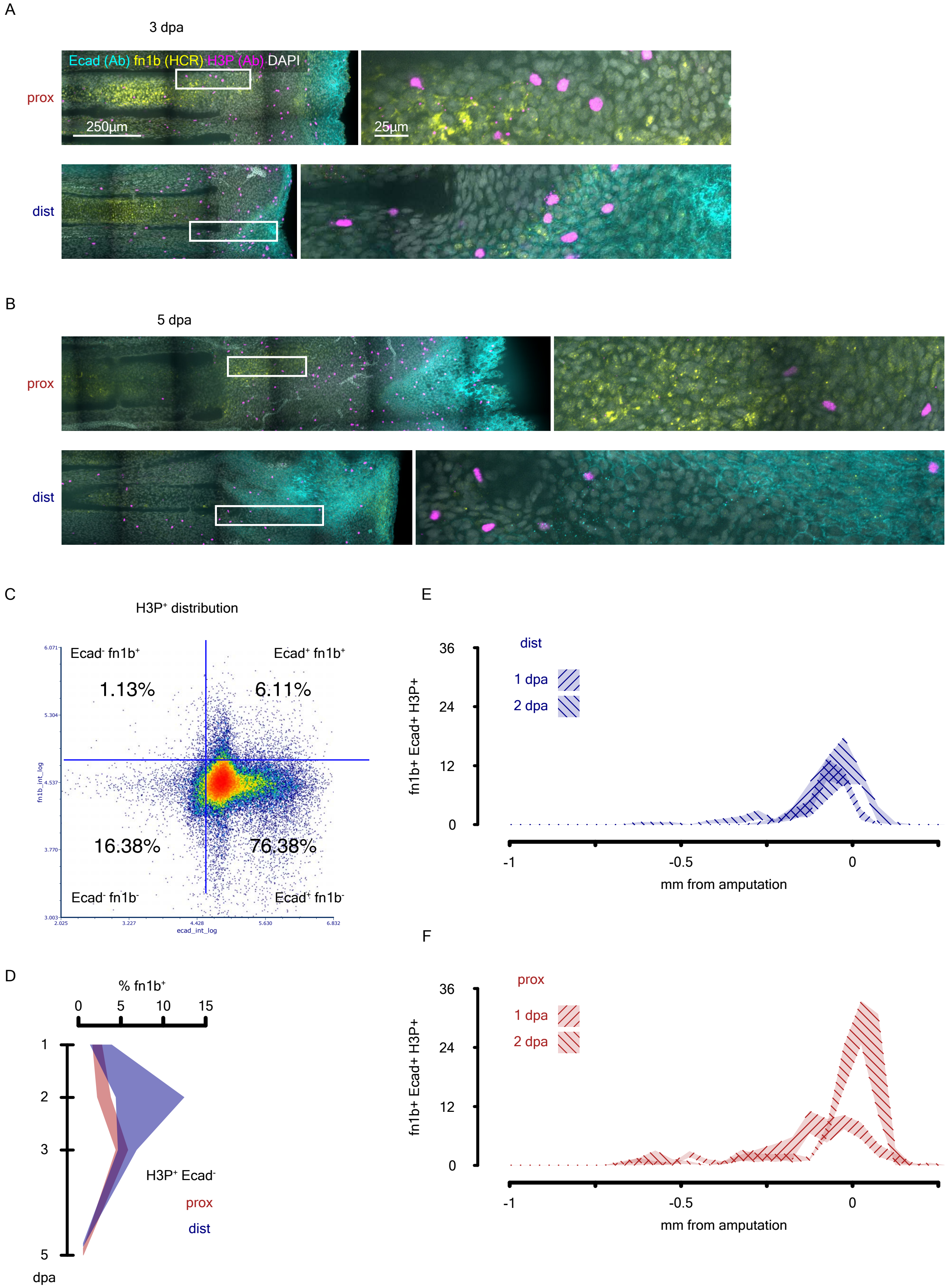

A
3 dpa
Ecad (Ab) fn1b (HCR) H3P (Ab) DAPI
prox
250µm
25µm
dist
B
5 dpa
prox
dist
C
H3P⁺ distribution
Ecad⁻ fn1b⁺
Ecad⁺ fn1b⁺
1.13%
6.11%
16.38%
76.38%
Ecad⁻ fn1b⁻
Ecad⁺ fn1b⁻
fn1b_int_log
ecad_int_log
D
% fn1b⁺
0
5
10
15
1
2
3
5
dpa
H3P⁺ Ecad⁻
prox
dist
E
dist
1 dpa
2 dpa
fn1b+ Ecad+ H3P+
36
24
12
0
-1
-0.5
0
mm from amputation
F
prox
1 dpa
2 dpa
fn1b+ Ecad+ H3P+
36
24
12
0
-1
-0.5
0
mm from amputation

**Supplementary Figure S4. *fn1b*$^{+}$ expression shuts down at later time points, *fn1b* and Ecad cytometry analysis, and spatial distribution of *fn1b*$^{+}$ Ecad$^{+}$ H3P$^{+}$ cells.**

(A) 3 dpa *fn1b* (HCR), Ecad (Ab), H3P (Ab) whole mount staining. (needs a scale bar)

(B) 5 dpa *fn1b* (HCR), Ecad (Ab), H3P (Ab) whole mount staining.

(C) Ecad and *fn1b* cytometry analysis on H3P$^{+}$ cells.

(D) % of *fn1b*$^{+}$ cells within the Ecad$^{-}$ H3P$^{+}$ population over time.

(E) Spatial distribution of *fn1b*$^{+}$ Ecad$^{+}$ H3P$^{+}$ cells in distal injuries at 1 and 2 dpa.

(F) Spatial distribution of *fn1b*$^{+}$ Ecad$^{+}$ H3P$^{+}$ cells in proximal injuries at 1 and 2 dpa.

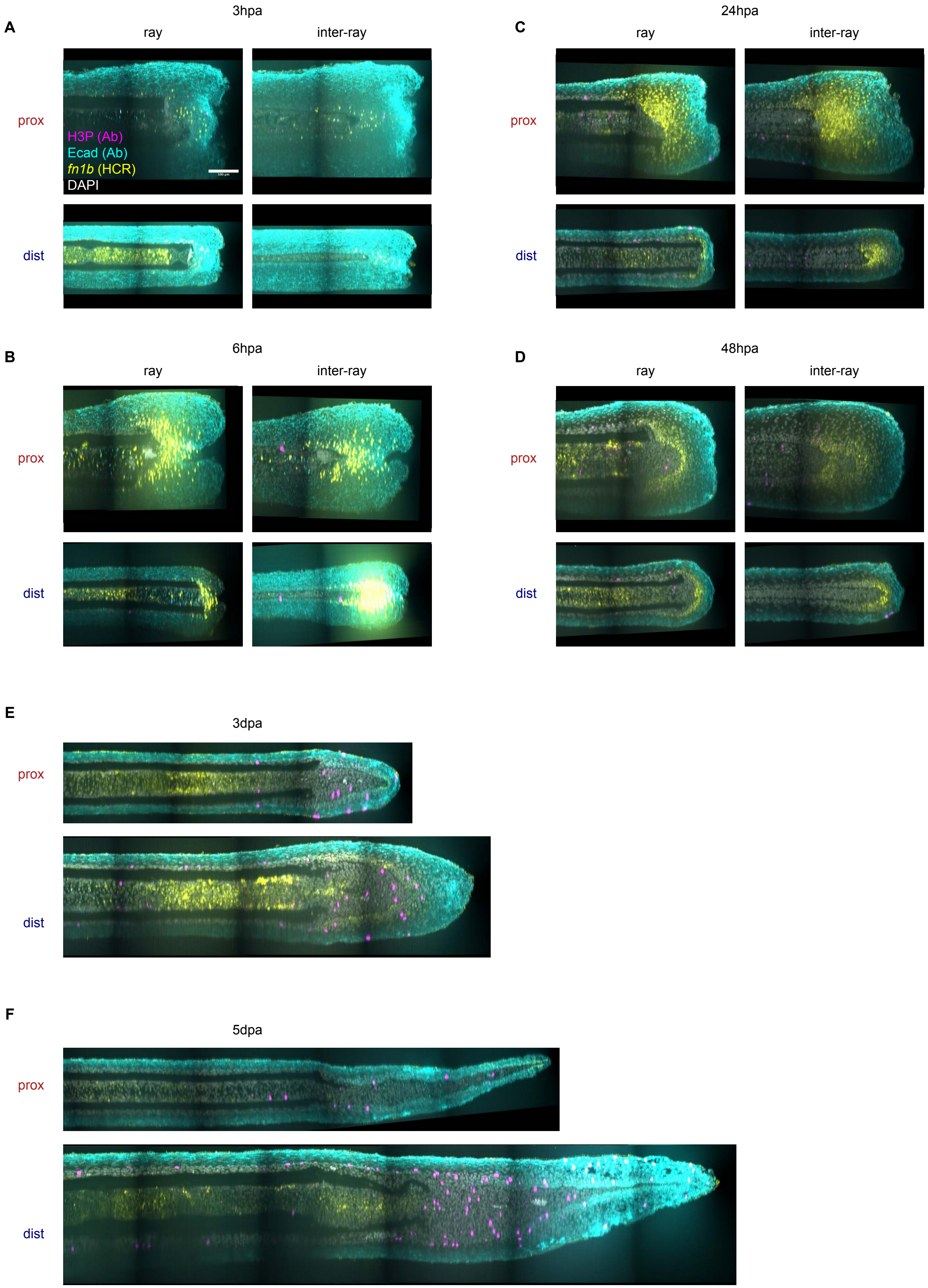
A
3hpa
ray
inter-ray
prox
H3P (Ab)
Ecad (Ab)
fn1b (HCR)
DAPI
dist
B
6hpa
ray
inter-ray
prox
dist
C
24hpa
ray
inter-ray
prox
dist
D
48hpa
ray
inter-ray
prox
dist
E
3dpa
prox
dist
F
5dpa
prox
dist

**Supplementary Figure S5. Orthogonal analysis of regeneration time course.**

(A) Ray and inter-ray orthogonal views of regenerating proximal and distal samples at 3 hpa *fn1b* (HCR), Ecad (Ab), H3P (Ab) whole mount staining. (fix scale bar)

(B) Same as in (A) at 6 hpa *fn1b* (HCR), Ecad (Ab), H3P (Ab) whole mount staining.

(C) Same as in (A) at 1 dpa *fn1b* (HCR), Ecad (Ab), H3P (Ab) whole mount staining.

(D) Same as in (A) at 2 dpa *fn1b* (HCR), Ecad (Ab), H3P (Ab) whole mount staining.

(E) Ray orthogonal views of regenerating proximal and distal samples at 3 dpa *fn1b* (HCR), Ecad (Ab), H3P (Ab) whole mount staining.

(F) Same as in (D) at 5 dpa *fn1b* (HCR), Ecad (Ab), H3P (Ab) whole mount staining.
